# Single-cell RNA-Sequencing Co-Expression Analysis with CFTR in Lung Tissue

**DOI:** 10.1101/2024.05.14.594146

**Authors:** Cheng Wang, Kayshani Kanagarajah, Amy Wong, Felix Ratjen, Lisa J Strug

## Abstract

1

**Background:** While cystic fibrosis is caused by loss-of-function variants in the *Cystic Fibrosis Transmembrane Conductance Regulator (CFTR)*, other modifier genes have been shown to associate with disease severity. Co-expression of modifiers with CFTR in normal tissue indicates a cooperative relationship and suggests the potential for compensation in the presence of CFTR dysfunction. We examined the co-expression relationships with CFTR in the lung using single cell RNA sequencing to pinpoint cell types and their modifiers involved in the forced expiratory volume in 1 second (FEV1)-based cystic fibrosis lung phenotype and support target cell-type prioritization for therapy ^1^.

**Methods:** SmartSeq2 single cell RNA sequencing data from non-cystic fibrosis lung tissue was used for evaluation of co-expression with CFTR and modifier genes. Zero-inflated negative binomial model was used to formally test the co-expression association. 10X Chromium based single cell RNA sequencing data from both cystic fibrosis and non-cystic fibrosis studies were assessed graphically to confirm conclusions from the SmartSeq2 primary analysis.

**Results:** Differentiating basal, club and alveolar epithelial type 2 cells were found to have high proportions of cells expressing CFTR as well as the greatest number of significant co-expression relationships with the modifiers. In particular, among alveolar epithelial type 2 cells, we observed a strong co-expression trio relationship between CFTR, SLC6A14 and SLC26A9 (p < 0.05).

**Conclusions:** CFTR-modifier gene co-expression suggests basal, club and alveolar epithelial type 2 cells show coordinated expression. Alveolar epithelial type 2 cells showed strong co-expression evidence with two of the most established cystic fibrosis modifier genes.

## 2 Introduction

Cystic fibrosis (CF) is a genetic disease caused by loss-of-function variants in the cystic fibrosis transmembrane conductance regulator (*CFTR*) ^2^, where progressive lung disease is a major contributor to morbidity and mortality ^3,4^. CF symptom severity varies among individuals with the same causal CFTR genotype and can be attributed in part to other genes, referred to as modifier genes ^5^.

Several CF modifier genes have been identified through genome-wide association studies ^6^. The mechanism by which these modifier genes contribute to CF in the presence of CFTR dysfunction is not entirely known although modifier genes such as *SLC9A3, SLC6A14* and *SLC26A9* encode alternative ion channels and transporters that regulate ion concentration and colocalize with CFTR at the apical membrane of epithelia ^7–9^. Modifier genes involved in immune response can alleviate CF symptoms from infection while other genes directly regulate the ciliary fluid layer ^10–12^.

CF paradigms for treatment have focused on CFTR correctors or potentiators with small molecules; alternative channels, transporters or pumps that can compensate for CFTR dysfunction; and gene correction through gene editing or replacement ^6^. Knowing the cell type to target is important for safety and efficacy concerns such as minimizing unwanted transgene expression in non-target cells ^13^.

Identifying potential cellular targets for the next generation of gene therapies requires convergent lines of evidence. Knowing where CFTR is expressed is necessary but likely not sufficient for this purpose as, for example, although ionocytes express a high proportion of CFTR they are likely not the major source of CFTR in the lungs ^14^. Previous study from He et al. showed extensive co-expression relationships among CFTR and CF modifier genes ^15^. As modifier genes have been linked to CF pathology across different affected organs, strong co-expression activity with CFTR itself in a given organ would support its role in CF and suggest relevant cell-types to target to ameliorate the clinical phenotype. Here, we hypothesize that CFTR and modifier genes of lung disease show co-expression in disease-relevant lung cell types, which can be investigated in future work as putative candidate cell-types for therapeutic target.

As demonstrated in bulk RNA-sequencing analysis using lung and nasal epithelium, CFTR and CF modifier genes are part of an active co-expression network in respiratory tissues ^15^. Recent applications of single cell RNA-sequencing (scRNA-Seq) technologies have enabled us to study CF co-expression networks at cellular resolution ^14,16^ and can provide information on the cellular target for the next generation of gene therapies. For the most comprehensive, unbiased investigation of important cell types for CF lung disease, the optimal study design would include all lung tissue and sequenced at high depth per cell. Single-cell sequencing of airways and lung parenchyma using the Smart-Seq2 technology has been completed ^17^ creating a valuable resource for co-expression analysis in lung tissue. Although the SS2 protocol is limited in the scope of cell types covered, the higher sequencing depth, when compared to the commonly used 10X Chromium protocol, allows for in-depth analysis of co-expression activities in the sampled cells with adequate power ^18^. This is sufficient for our goal of evaluating the cell type’s relevance to CF through studying CFTR co-expression rather than comprehensive characterization of airway cells.

In this study, we leveraged the Smart-Seq2 scRNA-Seq to study statistical significance of CFTR co-expression networks in various lung cell types and use the 10X data as replication support.

## 3 Materials and Methods

### 3.1 Study samples

The primary scRNA-seq resource used was obtained from Travaglini et al. ^17^. The donor cohort consisted of two males and one female individual aged from 46 to 75 years.

For replication and sensitivity analysis we also examined scRNA-Seq of lung tissue from non-CF and CF individuals sequenced using the 10X Chromium technology ^14,17,19^. The non-CF set included individuals in the Travaglini et al. study that were also 10X-sequenced and additional samples from the GTEx project (two males and one female individual aged 40 to 60 years) ^20^. Ten 10X-sequenced CF patients including six females, three males and one with unknown sex aged from 6 to 42 years old from ^14^ were also analyzed with most having at least one copy of the delta F508 *CFTR* mutation (n=8) (Table S1).

### 3.2 Single cell RNA-sequencing and data processing

Expression read count data were generated from SS2 scRNA-Seq, and detailed in Travaglini et al. ^17^. In addition to the filtering criteria applied by the original study, we also excluded 9,199 genes with no data in all cell samples. The final sample consisted of 9394 cells from 3 donors across 44 cell types (Table S2) with expression data from 49,484 genes (Table S3).

Unique molecular identifier (UMI) count data were obtained from the CF and healthy individual studies using 10X Chromium scRNA-Seq ^14,17,19^.

### 3.3 Statistical analysis

Co-expression relationships were evaluated using a zero-inflated negative binomial model with a log link function implemented in the R package pscl ^21,22^. To assess co-expression between two genes, read counts of one gene were regressed on that of the other gene and vice versa. For each cell, the proportion of genes with non-zero read counts was estimated to account for background effect on transcription detection in the model ^23^.

All model p-values were adjusted for multiple hypothesis testing using the FDR within cell type and summarized using the Cauchy combination method (R package ACAT) for each co-expression pair ^24–26^. All analysis was implemented in R ver. 3.5.1 ^27^. Functional relevance of the co-expressing genes was assessed using the GO term enrichment analysis tool Metascape ^28^.

Although tests for co-expression could not be implemented with the 10X design given corresponding power for co-expression detection, proportions of cells expressing both CFTR and a given gene in different cell types were counted, and used to determine whether there was replicated evidence for the Smart-Seq co-expression statistical analysis results.

### 3.4 Co-expression analysis

CFTR co-expression was analyzed with CF lung disease modifiers ^7,29–41^ and 157 genes of the apical plasma membrane constituents (Table S4 and S5) ^41^. Exploratory genome-wide co-expression analysis were also investigated.

## 4 Results

### 4.1 CFTR expression across different lung cell types

From the non-CF scRNA-Seq resource of Travaglini et al. ^17^ using the SS2 technology, the expression of *CFTR* was variable across different lung cell types (Figure 1). Twenty-five cell types had detectable CFTR expression (Table S6) and were included in the subsequent co-expression analysis.

**Figure 1.**
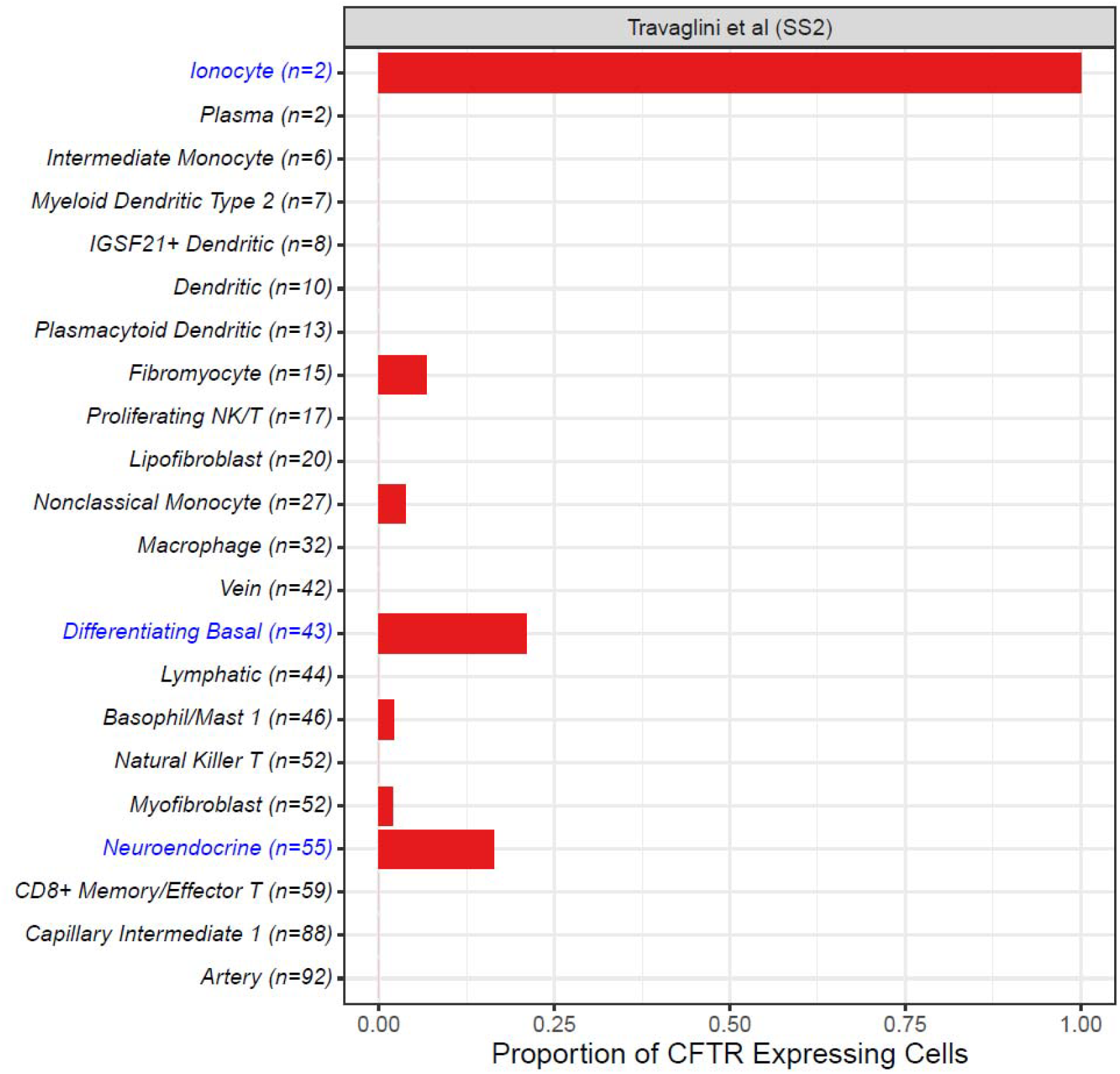
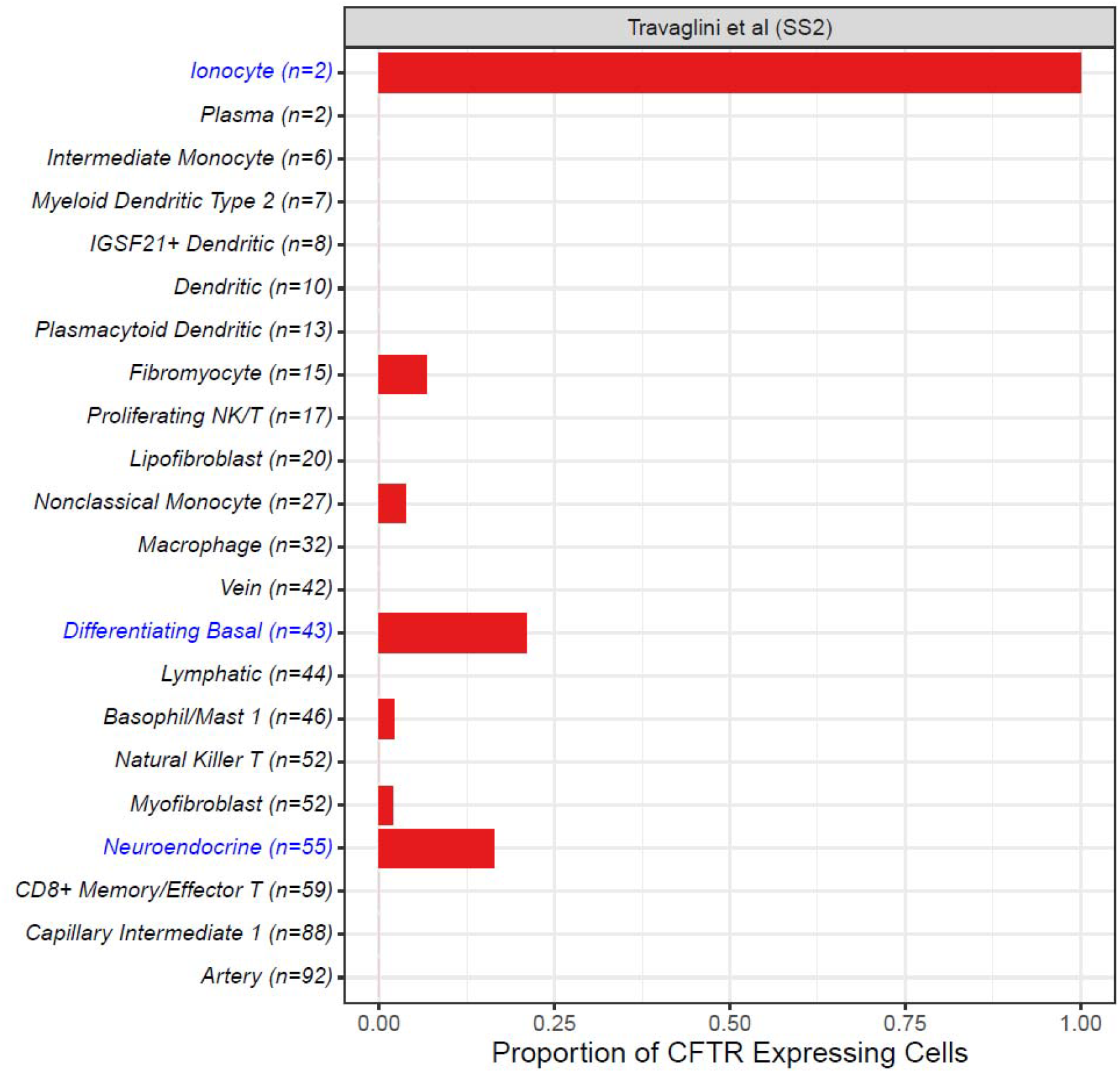
Proportion of cells expressing CFTR among lung cell types in the non-CF Smart-Seq2 data ^17^. Each bar presents the proportion of cells with detected CFTR expression. Cell types are arranged by their sample size in ascending order and denoted in the label. Ionocytes, goblets, alveolar epithelial type 2, club, signaling alveolar epithelial type 2, differentiating basal and neuroendocrine cells are highlighted in blue. SS2: Smart-Seq2.

The non-CF cell types with a high proportion of CFTR-expressing cells were ionocytes (100%), goblets (59%), alveolar epithelial type 2 (AT2) (41%), club (34%), signaling alveolar epithelial type 2 (sAT2) (34%), differentiating basal (21%) and neuroendocrine cells (16%).

Similar patterns of CFTR expression were also observed in the non-CF 10X Chromium data sets (Figure S1, S2). Among Travaglini et al. ^17^ 10X samples, high proportions of CFTR expressing cells were found in ionocytes, goblets, differentiating basal club, AT2 and sAT2 cell types. In the GTEx data, AT2, club and mast cells had the highest proportion of CFTR detected. In the CF 10X samples, ionocytes expressed CFTR in most of its cells, followed by secretory, FOXN4+ and basal cells (Figure S3).

### 4.2 CFTR is co-expressed with CF lung modifier and apical constituent genes in alveolar epithelial type 2 cells, differentiating basal and club cells

We examined patterns of CFTR co-expression with CF modifiers and apical genes (Table S4 and S5) in the SS2 and 10X Chromium data. For non-CF samples in the SS2 data, differentiating basal, AT2 and club cells demonstrated the greatest number of CFTR co-expressing genes across different lung cell types (Figure 2), indicating large co-expression networks. Ionocytes showed no significant co-expressing genes due to its limited sample size of n=2.

**Figure 2.**
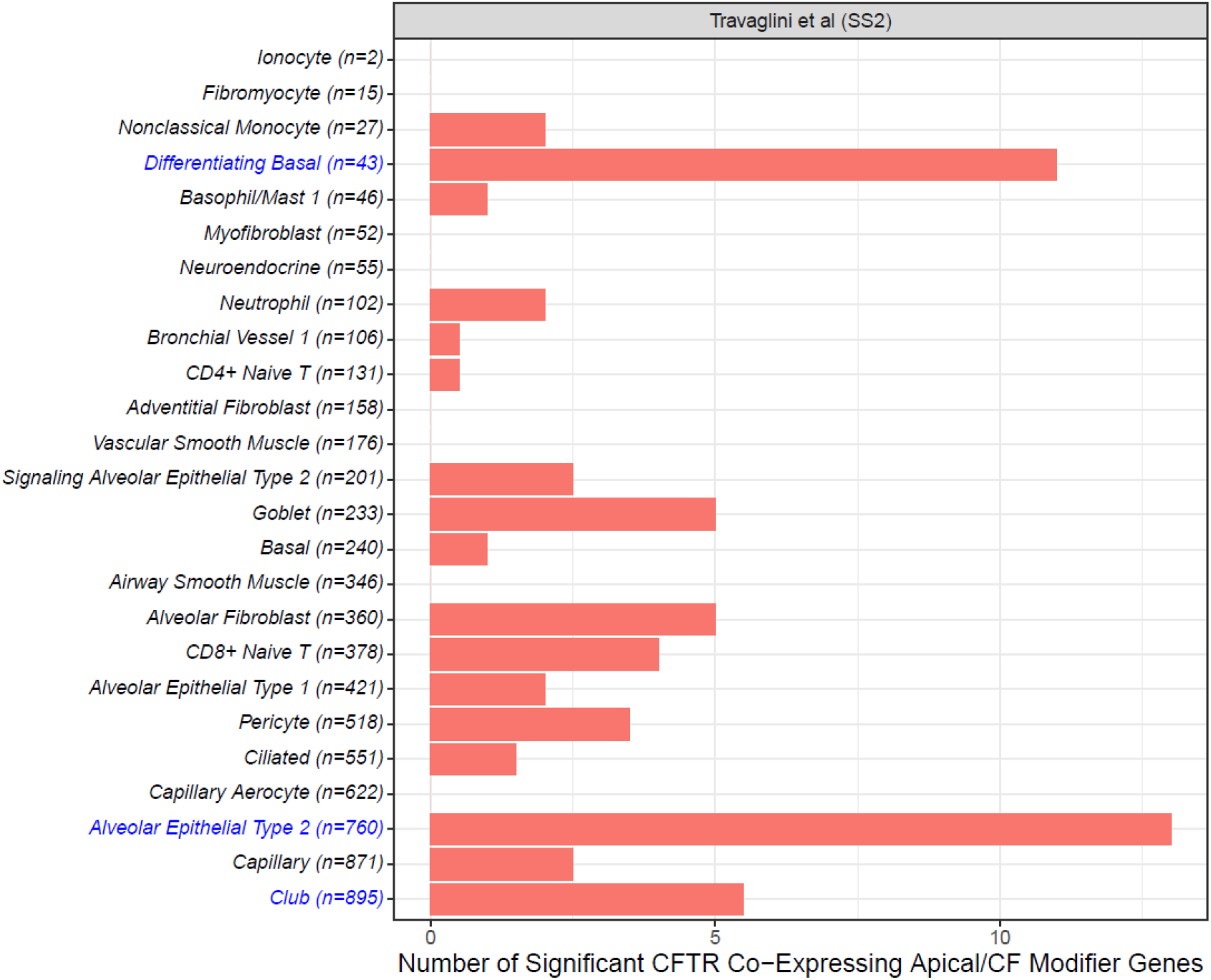
Number of significant co-expression relationships detected among cystic fibrosis modifiers including apical constituents (p < 0.05) across different lung cell types ordered by number of cells sequenced in the non-CF Smart-Seq2 sample ^17^. The contribution to the number of co-expression relationships for each cell type was based on the two models fitted for each gene, where each significant model result increased the count by 0.5.

Sample size for each cell type is denoted in the label. Alveolar epithelial type 2, club, and differentiating basal cells are highlighted in blue. SS2: Smart-Seq2.

Figure 3 summarizes the GO term enrichment test for cell types with any significant CFTR co-expressing gene. Biological regulation and response to stimulus were associated with AT2 and differentiating basal cells.

**Figure 3.**
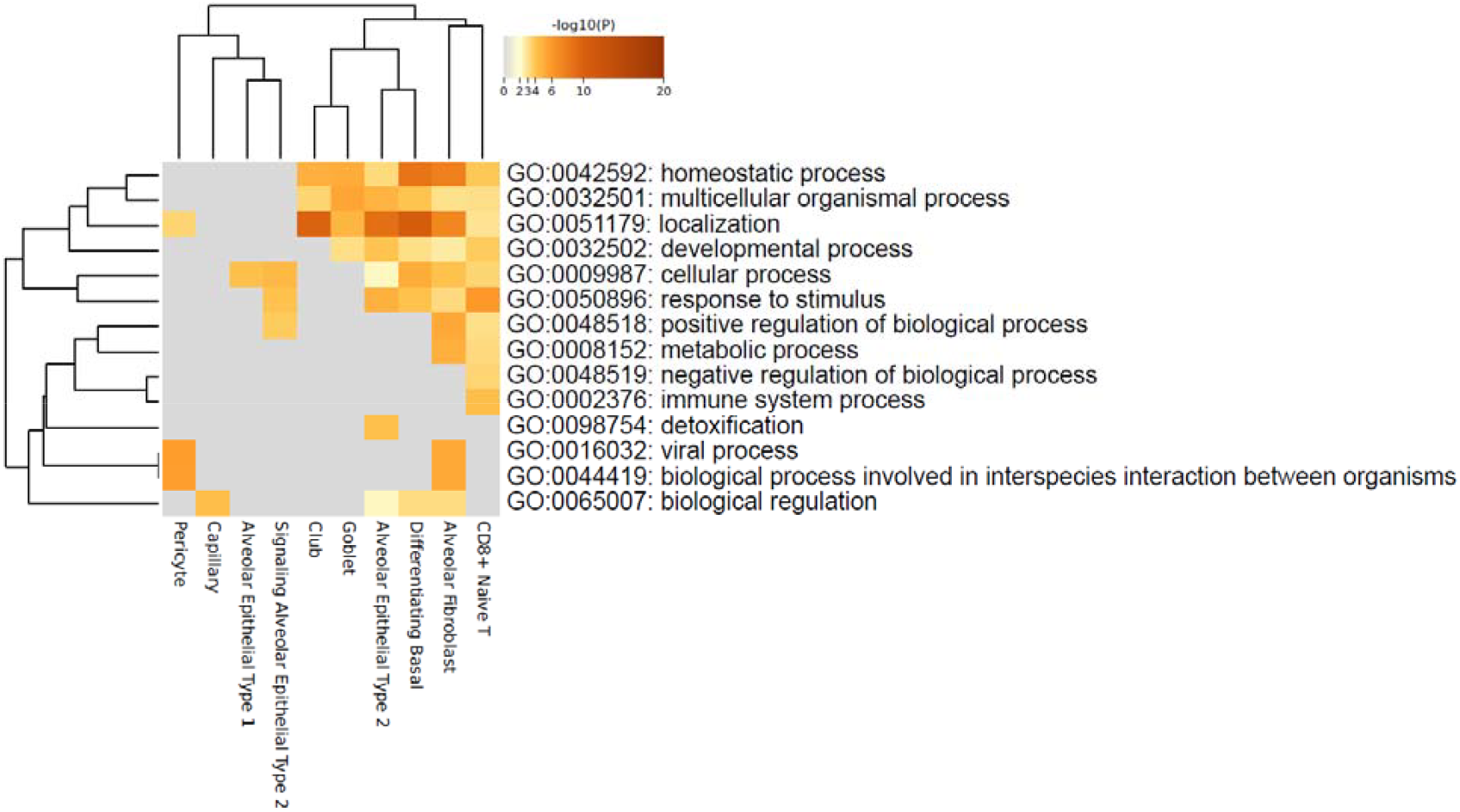
Functional GO term enrichment among the significant cystic fibrosis modifier and apical constituent genes that co-express with CFTR in the non-CF Smart-Seq2 sample ^17^. The heatmap displays the significance of association between a cell type’s o-expressing genes and a given GO term. Cell type (horizontal axis) and GO term (vertical axis) were clustered based on their enrichment test results. Localization is expected given the inclusion of 157 apical constituents defined by GO Annotation.

Among the 10x Chromium samples, we used the number of cells in a given cell type that had at least one gene and CFTR transcript as a proxy for a CFTR co-expressing gene to investigate support/replication of the SmartSeq co-expression results. Using this definition, the non-CF 10X sample from Travaglini et al. ^17^ also showed that AT2, differentiating basal and club cells had the highest number of co-expressing genes (Figure S4). The GTEx non-CF 10X sample also showed support for the significant co-expressing genes in AT2 and club cells (Figure S5). Although a different categorization of cell types was reported in the CF 10X sample, it demonstrated a high number of co-expressing genes for its broad basal cell type (Figure S6).

Similar patterns of CFTR co-expression were observed when we expanded the analysis to the whole genome in the SS2 and 10X Chromium data (Supplement section 2.3)

### 4.3 CFTR is in a co-expressing trio with SLC6A14 and SLC26A9 in alveolar epithelial type 2 cells

AT2, club and differentiating basal cells demonstrate significant co-expression relationships with CFTR. We examined these three cell types more closely by evaluating the mutual co-expression relationships among CFTR and CF modifiers in the non-CF SS2 samples (Figure 4 and S12).

AT2 cells showed a large number of significant co-expressing genes when investigating the CFTR-CF Modifier network, compared to club and differentiating basal cells.

**Figure 4.**
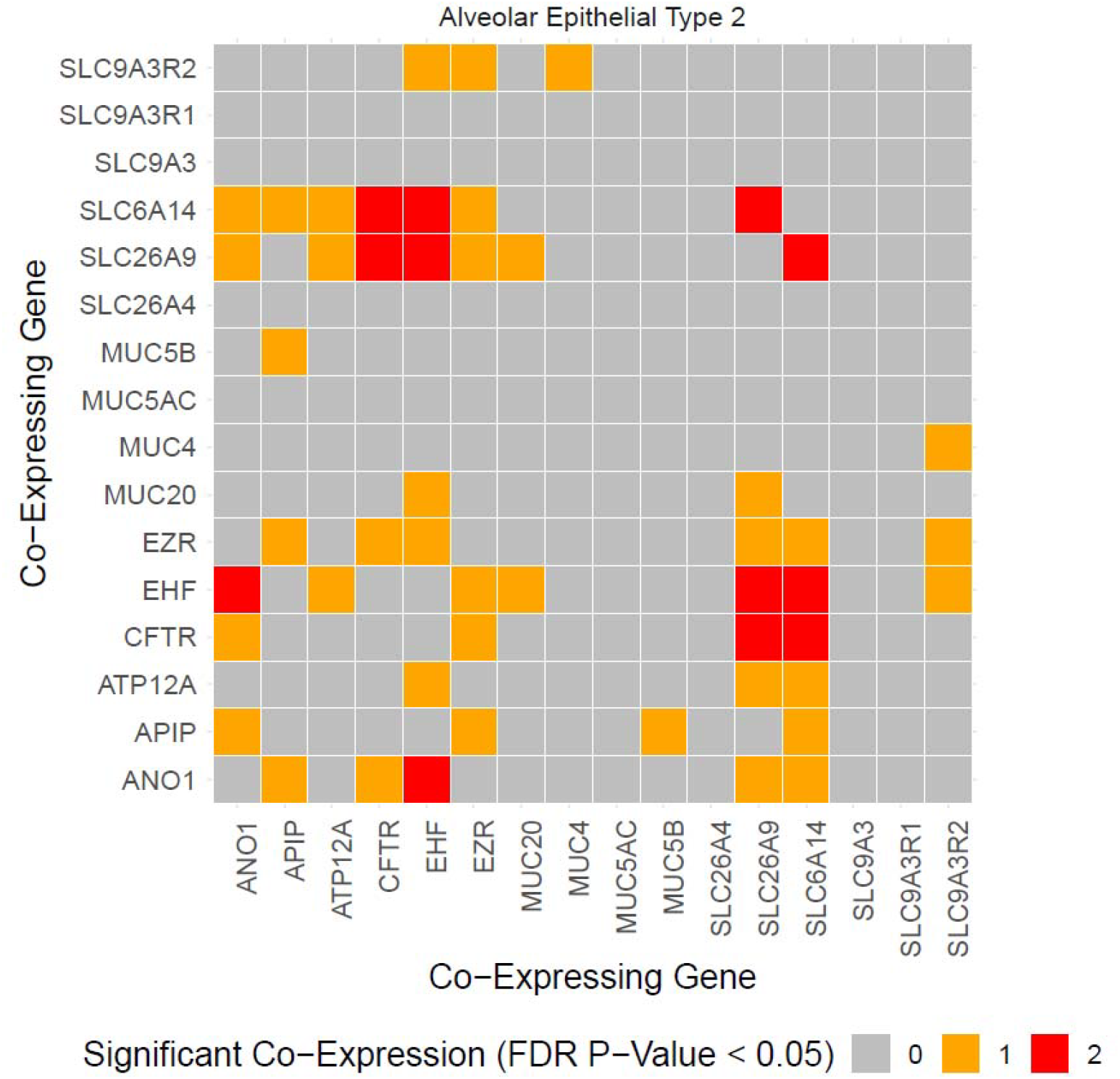
Co-expression relations among CFTR and cystic fibrosis modifier genes in alveolar epithelial type 2 from the non-CF Smart-Seq2 sample ^17^. Significant co-expression relationships are shown in red (both co-expression models significant) or yellow (one significant co-expression model) for each pair of genes.

In particular, CFTR was noted to have significant co-expression with *SLC6A14* and *SLC26A9* (Figure 4) and SLC6A14 and SLC26A9 were also co-expressed with each other in the same cell type (and with several other established CF lung modifier genes, although not with SLC9A3). This co-expression trio relationship was observed only in the AT2 cells, and remained significant after multiple hypothesis test corrections (Table 1). Expression of each of these three genes were also individually higher in the AT2 cells compared to other cell types (Figure S13).

**Table 1.** CFTR, SLC6A14 and SLC26A9 co-expression relationship in alveolar epithelial type 2 cells in the non-CF Smart-Seq2 sample ^17^.

| Cell Type | Outcome Gene | Predictor Gene | Effect Size | P-Value | FDR P-Value | Cauchy Combination P-Value |
| --- | --- | --- | --- | --- | --- | --- |
| Alveolar Epithelial Type 2 | CFTR | SLC6A14 | 0.00035 | $2.58 \times 10^{-36}$ | $3.30 \times 10^{-34}$ | $6.60 \times 10^{-34}$ |
| Alveolar Epithelial Type 2 | SLC6A14 | CFTR | 0.00162 | $9.41 \times 10^{-07}$ | $2.41 \times 10^{-05}$ | |
| Alveolar Epithelial Type 2 | CFTR | SLC26A9 | 0.00073 | $6.69 \times 10^{-06}$ | 0.00016 | $2.96 \times 10^{-4}$ |
| Alveolar Epithelial Type 2 | SLC26A9 | CFTR | 0.00059 | 0.00018 | 0.00296 |  |
| Alveolar Epithelial Type 2 | SLC6A14 | SLC26A9 | 0.00039 | 0.00192 | 0.01892 | $4.16 \times 10^{-10}$ |
| Alveolar Epithelial Type 2 | SLC26A9 | SLC6A14 | 0.00015 | $2.43 \times 10^{-12}$ | $2.08 \times 10^{-10}$ | |

Similar patterns of co-expression between these three genes were supported in the 10X Chromium data sets (Figure S14, S15). Both the non-CF 10X studies of Travaglini et al. ^17^ and GTEx ^20^ showed a high proportion of cells co-expressing CFTR-SLC26A9, CFTR-SLC6A14 and SLC26A9-SLC6A14 in AT2 cells compared to other cell types. The proportion of cells expressing each gene was higher in AT2 cells (Figure S17, S18).

Unfortunately, AT2 cells were not included in the CF 10X data (Figure S16), but the other cell types did not show a high proportion of co-expressing cells for CFTR-SLC26A9 and SLC26A9-SLC6A14 gene pairs. No cell type in the data showed high expression proportion for all three genes consistent with SS2 and other 10X data results (Figure S19).

## 5 Discussion

This study investigated the co-expression patterns of *CFTR* with CF-related genes in different lung cell types and found significant co-expression in AT2 cells. This was surprising to us since the acinar airways and alveoli are not presumed to be the sites of CF lung disease and as a consequence little attention has been paid to this cell type in CF research altogether ^42,43^.

Bulk RNA-sequencing analysis showed a high number of co-expression relationships among CFTR and CF modifier genes ^15^. When examining the number of active co-expressing genes, differentiating basal, AT2 and club cells have the greatest number of CF modifier genes co-expressed with CFTR, aligning with the results from the bulk RNA-sequencing studies ^15^. Studies have shown that in addition to CFTR, modifier genes also contribute to the lung disease severity phenotype in CF ^6,7,41^. Hence, considering CFTR’s co-expressing relationships with modifier genes can assist in narrowing down the important and relevant cell types in the lung for therapeutic target and functional study.

Among the CF modifiers, He et al showed in a human nasal and bronchial tissue bulk RNA-sequencing study that SLC6A14 is one of the most active co-expressing gene, with respect to both CFTR and other modifiers ^15^. Here, we were able to detect significant co-expression with two of the most widely investigated modifier genes in CF, *SLC26A9* and *SLC6A14* ^44–49^, with each other and with *CFTR* in AT2 cells. The expression of these two genes and CFTR are highest in AT2, sAT2, club, differentiating basal and goblet cells compared to the other cell types (Figure S13). Relatively high co-expression in these cell types is further supported in the independent 10X data sets (Figure S14 – S16).

*SLC26A9* is a decoupled chloride transporter that has been shown to interact with CFTR ^48,50^ and modify response to CFTR modulator therapies ^9,32^. Whereas *SLC6A14*, identified as a lung disease modifier in the largest GWAS of CF lung disease to date, has been shown to modulate response to Pseudomonas aeruginosa infection in the airway ^45^, a common CF respiratory infection that contributes to cycles of infection and inflammation and progressive lung disease, and CFTR channel function ^51^. Co-expression of CFTR with these two important CF modifier genes suggests that AT2 cells could play a role in CF that requires further investigation. In addition to AT2 cells, other significant co-expression relationships involve SLC9A3R2 in the differentiating basal cells (CFTR-SLC9A3R2) and the club cells (MUC4-SLC9A3R2) (Supplement section 2.4).

The significant, positive linear co-expression of CFTR, SLC26A9 and SLC6A14 in AT2 cells and that SLC6A14 and SLC26A9 are CF lung disease modifier genes, posits that disruption of CFTR in CF could disrupt the balance of these transporters and provide one mechanism and site of action by which the modifiers contribute to lung disease severity. In mice the SLC26A9 knockout led to early mortality due to pulmonary complications and reduced airspace in the lungs ^52^, attributed to AT2 localization.

Despite strong, replicated support in this study for co-expression of CFTR and modifiers in AT2 cells, basal and club cells, there are several limitations that should be noted. The sequencing depth of scRNA-seq technology is lower compared to bulk RNA-seq. As a result, detection of genes with low expression is more difficult due to dropout. Our primary analysis data were generated using the SS2 single cell library preparation protocol. This procedure allows for deeper sequencing depth compared to the more commonly used 10X Chromium protocols. One caveat of using SS2 is that it counts the number of aligned sequencing reads for expression measures instead of unique molecular identifier. Thus, it was reassuring that when we compared our main results with an analysis performed using the 10X Chromium data sets, the 10X result was supportive of our main conclusions.

The comprehensiveness of our study is also limited due several factors. Cell types with appreciable CFTR expression but low count such as the ionocytes could not be analyzed for co-expression. Samples for some cell types including the differentiating basal cells came from a single individual, which could potentially reduce the generalizability of our findings. Similarly, sampling of tissues at limited locations in the lung airway can affect the results as well. In addition to sampling limitations, using co-expression activity to infer cell type’s relevance to CF would miss other cellular mechanism for CF involvement. For example, compensatory or regulatory actions on CFTR may originate in neighboring cell types rather than the CFTR-expressing cell itself. This would not be fully reflected in the CFTR co-expression activities. There are also likely other cell types where CFTR are co-expressed with modifier genes but presumably to a lesser degree that was not detectable with the present available resources due to sequencing dropout or low cell samples available. Thus, the absence of CFTR co-expression in our results should not be interpreted against the relevance of the other cell types in CF.

In conclusion, our study explored the co-expression networks of CFTR in different lung cell types, with emphasis on modifier genes under the presumption that co-expression of CFTR and modifiers could pinpoint CF lung disease relevant cell types. Surprisingly, we observed strong CFTR and lung modifier gene co-expression in AT2 cells, suggesting there may be a previously unappreciated role for AT2 cells in CF and the alveolar compartment of the lung that should be considered for further study.

## Supporting information

Methods and Results Supplement

## Acknowledgements

This research was supported by funding from the Canada Research Chairs Program; the Canadian Institutes of Health Research (CIHR) Foundation Grant, FRN: 167282; and a peer-reviewed Cystic Fibrosis (CF) Canada 2022 Clinical Research Grant jointly funded by CF Canada and Canadian Institutes of Health Research Institute of Circulatory and Respiratory Health (CIHR-ICRH), FRN: BCG 187014.

