## Supplementary material for "Single-cell RNA-Sequencing Co-Expression Analysis with CFTR in Lung Tissue": Methods and Results Supplement

Correspondance to:

Lisa J Strug

The Centre for Applied Genomics and

Program in Genetics and Genome Biology

The Hospital for Sick Children Research Institute

686 Bay Street, Room 12-9705, Toronto, ON, Canada

M5G 0A4

### 1 Methods Supplement

Table S1. CFTR mutation genotypes for the patients included in the CF 10X Chromium data from Carraro et al. ^1^.

| **Patient** | **CFTR Genotype 1** | **CFTR Genotype 2** |
| --- | --- | --- |
| 1 | ΔF508 | ΔF508 |
| 2 | ΔF508 | ΔF508 |
| 3 | ΔF508 | ΔF508 |
| 4 | ΔF508 | ΔF508 |
| 5 | ΔF508 | 1078delT |
| 6 | ΔF508 | 1811+1.6kbA>G |
| 7 | ΔF508 | R553X |
| 8 | ΔF508 | G551D |
| 9 | G542x | L1077p |
| 10 | G542x | R785x |

Table S2. List of cell samples in the in the non-CF Smart-Seq2 data ^2^.

| **Cell Type** | **Donors** | **Cells** |
| --- | --- | --- |
| Adventitial Fibroblast | 3 | 158 |
| Airway Smooth Muscle | 3 | 346 |
| Alveolar Epithelial Type 1 | 3 | 421 |
| Alveolar Epithelial Type 2 | 3 | 760 |
| Alveolar Fibroblast | 3 | 360 |
| Artery | 3 | 92 |
| B | 3 | 129 |
| Basal | 3 | 240 |
| Basophil/Mast 1 | 3 | 46 |
| Bronchial Vessel 1 | 1 | 106 |
| Capillary | 3 | 871 |
| Capillary Aerocyte | 3 | 622 |
| Capillary Intermediate 1 | 1 | 88 |
| CD4+ Memory/Effector T | 1 | 126 |
| CD4+ Naive T | 2 | 131 |
| CD8+ Memory/Effector T | 1 | 59 |
| CD8+ Naive T | 2 | 378 |
| Ciliated | 3 | 551 |
| Classical Monocyte | 1 | 106 |
| Club | 3 | 895 |
| Dendritic | 1 | 10 |
| Differentiating Basal | 1 | 43 |
| Fibromyocyte | 1 | 15 |
| Goblet | 3 | 233 |
| IGSF21+ Dendritic | 2 | 8 |
| Intermediate Monocyte | 2 | 6 |
| Ionocyte | 1 | 2 |
| Lipofibroblast | 1 | 20 |
| Lymphatic | 3 | 44 |
| Macrophage | 2 | 32 |
| Myeloid Dendritic Type 2 | 1 | 7 |
| Myofibroblast | 2 | 52 |
| Natural Killer | 3 | 1232 |
| Natural Killer T | 2 | 52 |
| Neuroendocrine | 2 | 55 |
| Neutrophil | 3 | 102 |
| Nonclassical Monocyte | 2 | 27 |
| Pericyte | 3 | 518 |
| Plasma | 1 | 2 |
| Plasmacytoid Dendritic | 3 | 13 |
| Proliferating NK/T | 2 | 17 |
| Signaling Alveolar Epithelial Type 2 | 2 | 201 |
| Vascular Smooth Muscle | 3 | 176 |
| Vein | 1 | 42 |

Table S3. Type of genes found in the non-CF Smart-Seq2 data ^2^ according to GENCODE version 29 annotation.

| **Gene Type** | **Number of Genes** |
| --- | --- |
| Protein Coding | 18664 |
| 3prime Overlapping ncRNA | 12 |
| Antisense | 3936 |
| Bidirectional Promoter lncRNA | 63 |
| IG C Gene | 13 |
| IG C Pseudogene | 6 |
| IG J Gene | 8 |
| IG V Gene | 2 |
| IG V Pseudogene | 2 |
| lincRNA | 6599 |
| Macro lncRNA | 1 |
| miRNA | 906 |
| Misc RNA | 1755 |
| Polymorphic Pseudogene | 33 |
| Processed Pseudogene | 8421 |
| Processed Transcript | 266 |
| Pseudogene | 15 |
| Ribozyme | 3 |
| rRNA | 17 |
| rRNA Pseudogene | 298 |
| scaRNA | 33 |
| scRNA | 1 |
| Sense Intronic | 722 |
| Sense Overlapping | 157 |
| snoRNA | 619 |
| snRNA | 127 |
| sRNA | 4 |
| TEC | 942 |
| Transcribed Processed Pseudogene | 395 |
| Transcribed Unitary Pseudogene | 112 |
| Transcribed Unprocessed Pseudogene | 773 |
| Translated Processed Pseudogene | 2 |
| TR C Gene | 6 |
| TR D Gene | 3 |
| TR J Gene | 64 |
| TR J Pseudogene | 4 |
| TR V Gene | 49 |
| TR V Pseudogene | 16 |
| Unitary Pseudogene | 76 |
| Unprocessed Pseudogene | 1820 |
| No Annotation Available | 2539 |

Table S4. List of Published CF modifier genes examined

| **Gene** | **Evidence** |
| --- | --- |
| ANO1 | Alternative chloride channel proposed as a potential therapeutic target for CF ^3^. Experimental evidence indicating co-expression with CFTR in basal cells as well as regulatory role in basal differentiation, which is disrupted in CF ^4–6^. |
| APIP | Involved in inflammation, apoptosis and methionine salvage pathway ^7^. GWAS evidence associated with CF lung disease severity ^8^. |
| ATP12A | Regulate pH balance via proton secretion with higher expression in CF airways ^9^. GWAS evidence associated with meconium ileus in CF ^10^. |
| EHF | Transcription factor involved in wound repair and repression of CFTR expression ^11^. GWAS evidence associated with CF lung disease severity ^8,12^. |
| MUC4, MUC20 | Prevent mucus from entering periciliary space and help with mucoiliary host defense. GWAS evidence associated with CF lung disease severity ^8^. |
| MUC5AC, MUC5B | Component of airway mucus and altered secretion in CF ^13^. |
| SLC26A4 | Regulate bicarbonate concentration and interact with CFTR ^14^. |
| SLC26A9 | Anion channel interacting with CFTR ^15,16^. GWAS evidence associated with meconium ileus, diabetes in CF, lung function and lung function response to CFTR-directed therapeutics ^17–20^. |
| SLC6A14 | Transport neutral and cationic amino acid in sodium and chloride dependent fashion. GWAS evidence associated with lung and meconium ileus disease in CF ^8,19^. |
| SLC9A3 | Regulate pH balance via exchanging Na+/H+ ions. GWAS evidence associated with lung and meconium ileus disease in CF ^8,10,19,21^. |
| SLC9A3R1, SLC9A3R2, EZR | Regulate SLC9A3 and CFTR ^22^. Association evidence with CF lung disease severity when in complex with SLC9A3 ^10,18,21^. |

CF: cystic fibrosis; GWAS: genome-wide association study

Table S5. List of apical plasma membrane constituent genes examined in the analysis

| ABCA7 | CD300LG | KCNE1 | P2RY4 | SLC22A5 | TGFBR1 |
| --- | --- | --- | --- | --- | --- |
| ABCB4 | CD44 | KCNMA1 | P2RY6 | SLC23A1 | TLR9 |
| ABCC2 | CDHR2 | KIAA1919 | PDPN | SLC23A2 | TRPM6 |
| ABCC6 | CFTR | KNCN | PFKM | SLC26A3 | TRPV5 |
| ABCG5 | CIB1 | LCT | PKHD1 | SLC26A4 | UMOD |
| ABCG8 | CLCA4 | LHFPL5 | PLB1 | SLC26A9 | UPK3A |
| ACY3 | CRB1 | LMO7 | PRKCI | SLC29A4 | VANGL2 |
| ADAM17 | CRB3 | LRP2 | PROM1 | SLC30A5 |  |
| ADRB2 | CSPG4 | LZTS1 | PROM2 | SLC34A2 |  |
| AJAP1 | CTSB | MAL | PTK2 | SLC34A3 |  |
| AKAP7 | CUBN | MAL2 | RAB14 | SLC39A4 |  |
| AKR1A1 | DPEP1 | MGAM | RAB27A | SLC3A2 |  |
| ANK2 | DPP4 | MIP | RHCG | SLC46A1 |  |
| ANXA6 | DUOX1 | MPDZ | S100G | SLC4A5 |  |
| AQP1 | DUOX2 | MREG | SCNN1A | SLC4A7 |  |
| AQP2 | EGFR | MSN | SCNN1B | SLC5A1 |  |
| AQP5 | ENPEP | MUC1 | SCNN1G | SLC5A12 |  |
| AQP8 | EPCAM | MUC20 | SHROOM2 | SLC5A8 |  |
| ATP1B1 | ERBB2 | MUC3A | SHROOM3 | SLC6A20 |  |
| ATP2B2 | ERBB3 | MUC3B | SHROOM4 | SLC7A5 |  |
| ATP4A | EZR | MYO1A | SI | SLC9A3 |  |
| ATP6V0A4 | F2RL2 | MYO7B | SLC10A2 | SLC9A3R1 |  |
| ATP6V0D1 | GIF | NAALADL1 | SLC11A2 | SLC9A3R2 |  |
| ATP6V0D2 | GJB6 | NPC1L1 | SLC12A2 | SLC9A4 |  |
| ATP6V1B1 | GNAT3 | NRG1 | SLC12A3 | STK39 |  |
| ATP6V1E1 | GPR64 | OTOA | SLC14A2 | STX3 |  |
| ATP8B1 | IGFBP2 | OTOG | SLC22A11 | STX4 |  |
| CA4 | IL6R | OXTR | SLC22A12 | STXBP3 |  |
| CACNB3 | INADL | P2RY1 | SLC22A18 | TCIRG1 |  |
| CAV1 | KCNA1 | P2RY2 | SLC22A4 | TF |  |

### 2 Results Supplement

#### 2.1 CFTR expression across different lung cell types

Table S6. List of cell types with detectable CFTR expression in the non-CF Smart-Seq2 data ^2^

| **Cell Type** | **Donors** | **Cells** | **Proportion of CFTR-Expressing Cell** |
| --- | --- | --- | --- |
| Ionocyte | 1 | 2 | 1 |
| Goblet | 3 | 233 | 0.587983 |
| Alveolar Epithelial Type 2 | 3 | 760 | 0.411842 |
| Club | 3 | 895 | 0.338547 |
| Signaling Alveolar Epithelial Type 2 | 2 | 201 | 0.338308 |
| Differentiating Basal | 1 | 43 | 0.209302 |
| Neuroendocrine | 2 | 55 | 0.163636 |
| Fibromyocyte | 1 | 15 | 0.066667 |
| Basal | 3 | 240 | 0.0625 |
| Nonclassical Monocyte | 2 | 27 | 0.037037 |
| Basophil/Mast 1 | 3 | 46 | 0.021739 |
| Myofibroblast | 2 | 52 | 0.019231 |
| Bronchial Vessel 1 | 1 | 106 | 0.018868 |
| Ciliated | 3 | 551 | 0.018149 |
| CD4+ Naive T | 2 | 131 | 0.015267 |
| Alveolar Fibroblast | 3 | 360 | 0.011111 |
| Neutrophil | 3 | 102 | 0.009804 |
| Alveolar Epithelial Type 1 | 3 | 421 | 0.009501 |
| CD8+ Naive T | 2 | 378 | 0.007937 |
| Adventitial Fibroblast | 3 | 158 | 0.006329 |
| Pericyte | 3 | 518 | 0.005792 |
| Airway Smooth Muscle | 3 | 346 | 0.00578 |
| Vascular Smooth Muscle | 3 | 176 | 0.005682 |
| Capillary | 3 | 871 | 0.003444 |
| Capillary Aerocyte | 3 | 622 | 0.001608 |

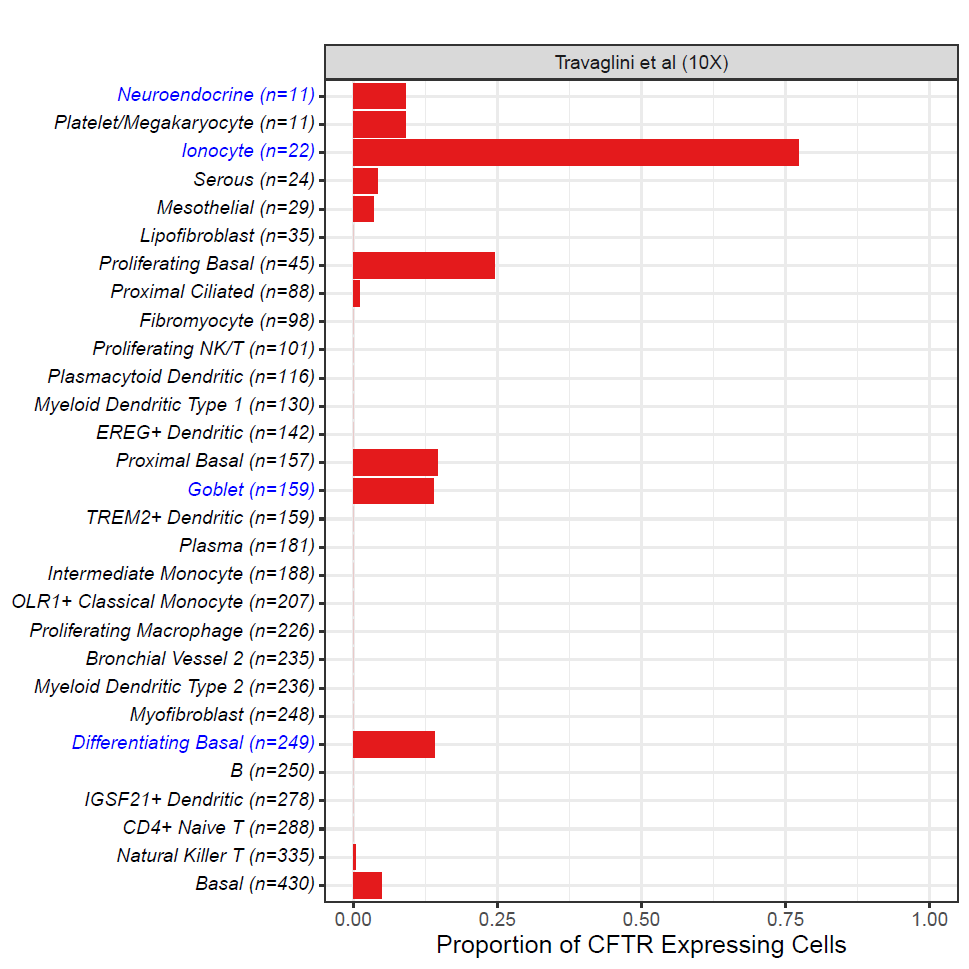

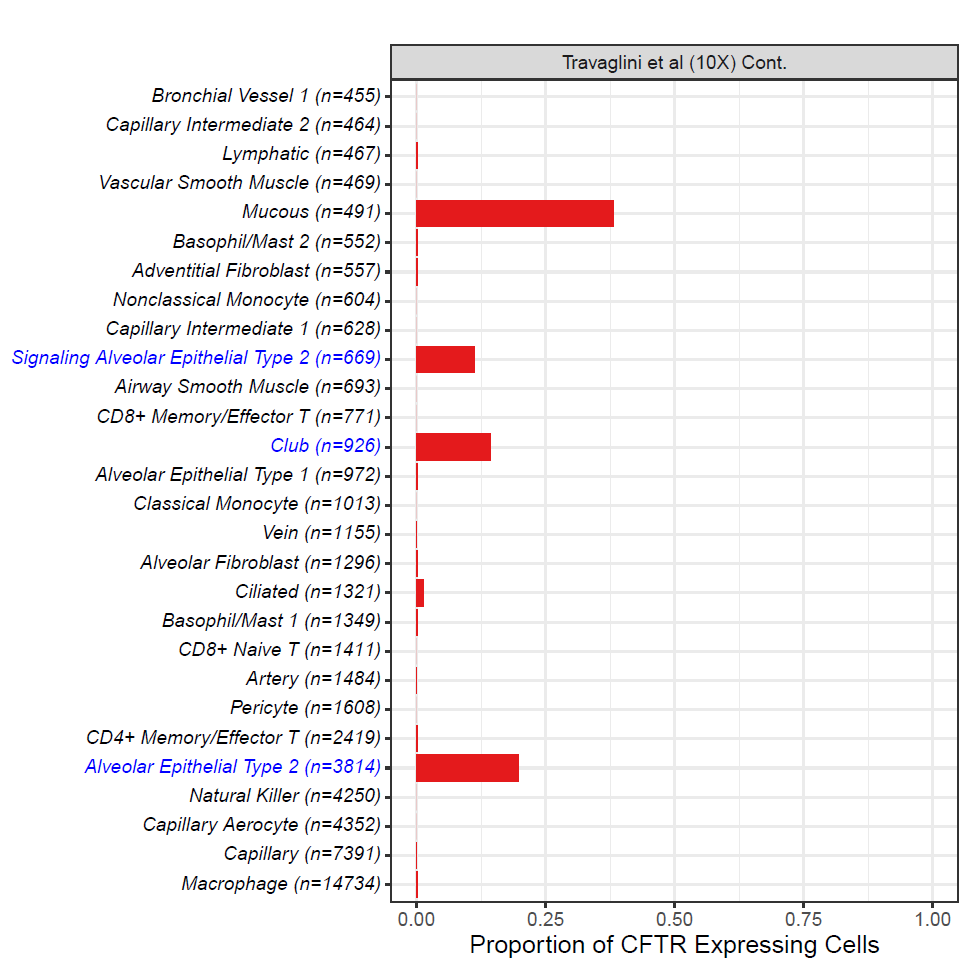

Figure S1. Proportion of samples expressing CFTR among lung cell types in the non-CF 10X Chromium data from Travaglini et al. ^2^.

Each bar presents the proportion of cells with detected CFTR expression. Cell types are arranged by their sample size in ascending order and denoted in the label. Ionocytes, goblets, alveolar epithelial type 2, club, signaling alveolar epithelial type 2, differentiating basal and neuroendocrine cells are highlighted in blue. 10X: 10X Chromium.

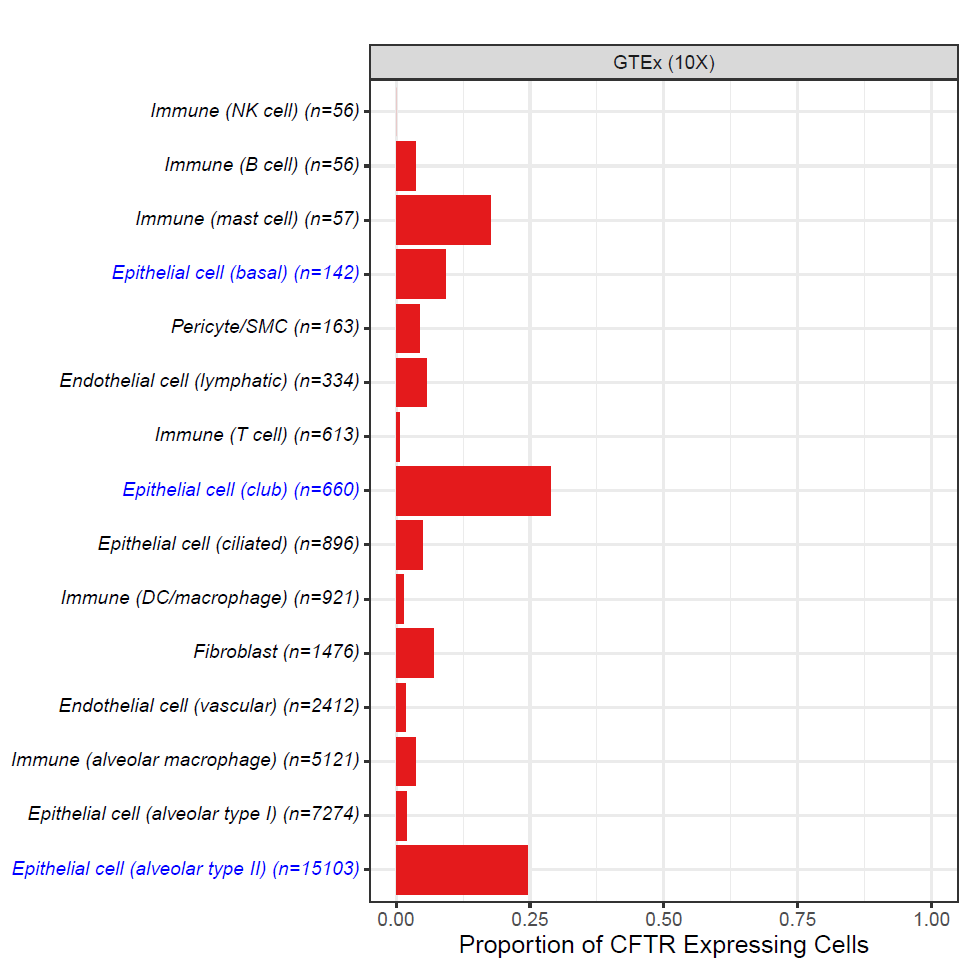

Figure S2. Proportion of samples expressing CFTR among lung cell types in the non-CF GTEx 10X Chromium data from GTEx ^23^.

Each bar presents the proportion of cells with detected CFTR expression. Cell types are arranged by their sample size in ascending order and denoted in the label. Alveolar epithelial type 2, club, and basal cells are highlighted in blue. 10X: 10X Chromium.

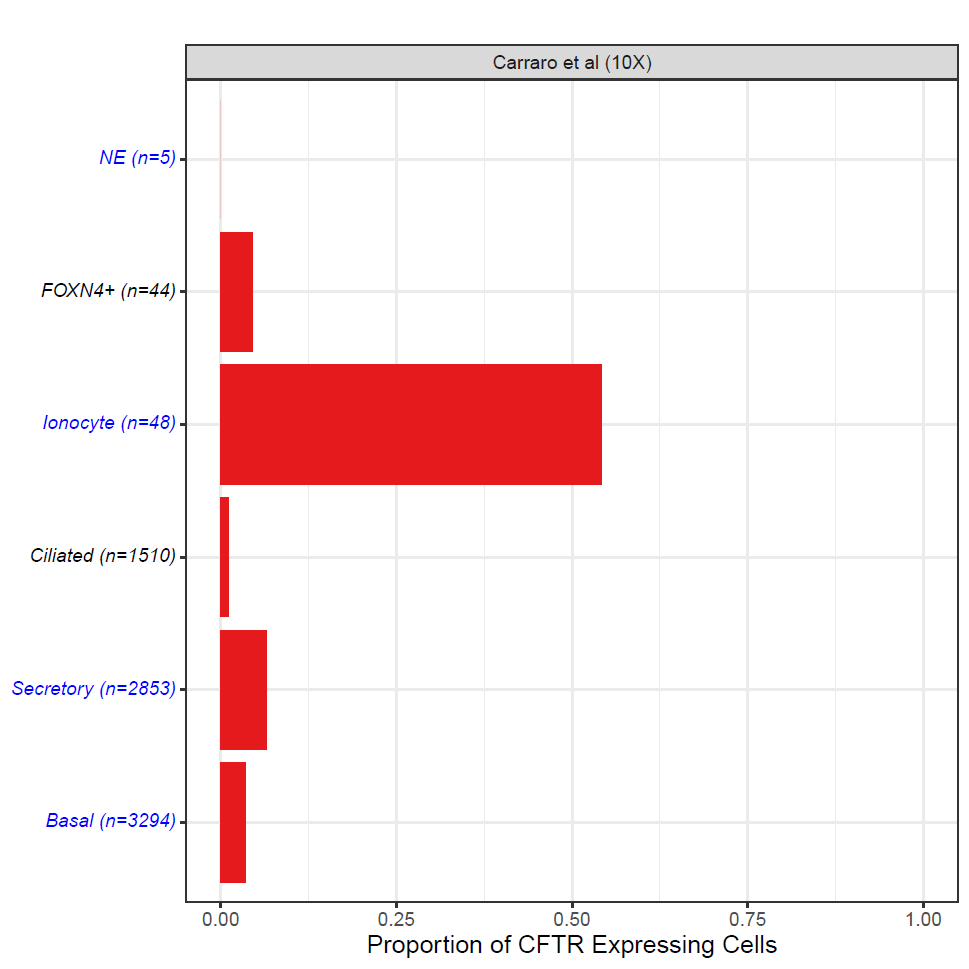

Figure S3. Proportion of samples expressing CFTR among lung cell types in the CF 10X Chromium data from Carraro et al. ^1^.

Each bar presents the proportion of cells with detected CFTR expression. Cell types are arranged by their sample size in ascending order and denoted in the label. Ionocytes, secretory, basal and neuroendocrine cells are highlighted in blue. NE: neuroendocrine; 10X: 10X Chromium.

#### 2.2 CFTR is co-expressed with CF lung modifier and apical constituent genes in alveolar epithelial type 2 cells, differentiating basal and club cells

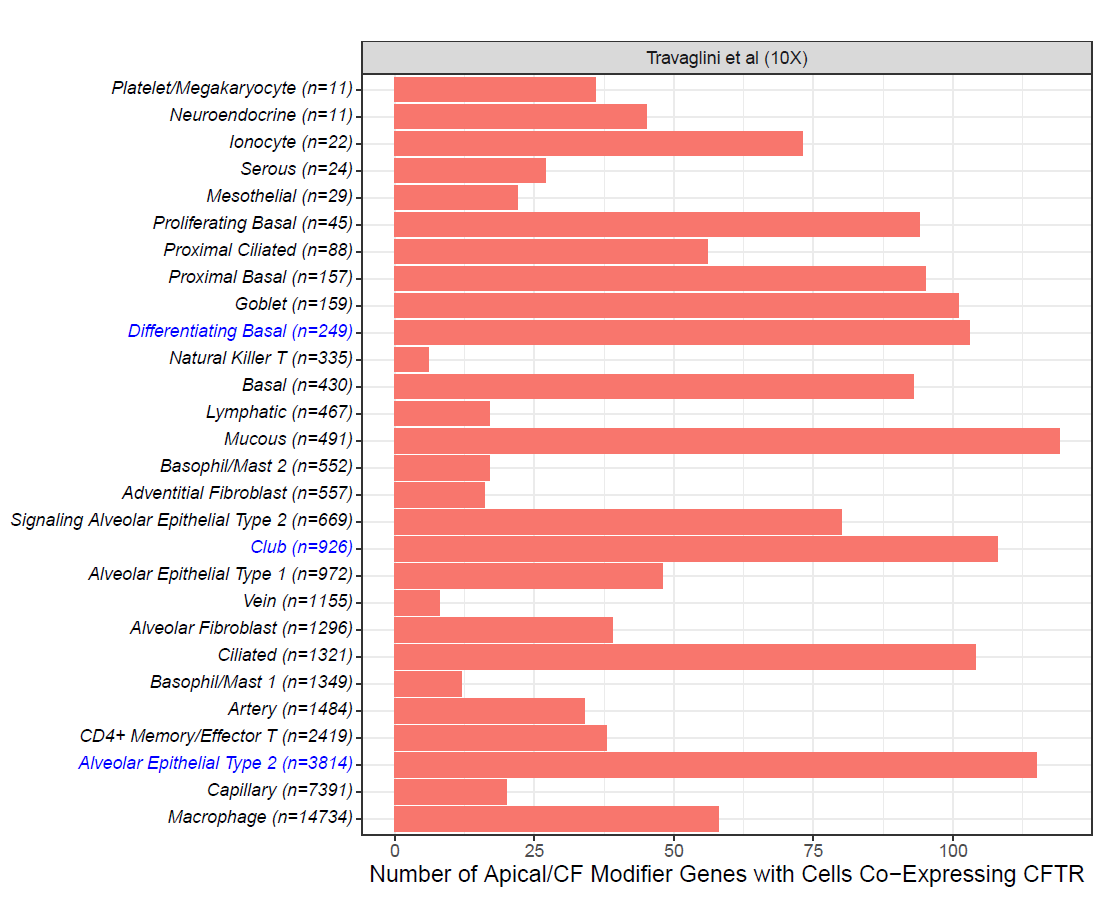

Figure S4. Number of cystic fibrosis modifier and apical constituent genes with at least one CFTR co-expressing cell across different lung cell type cells ordered by number of cells sequenced in the non-CF 10X Chromium data from Travaglini et al. ^2^.

A co-expressing cell was determined by observing detectable expression for CFTR and a given gene in the same cell. Sample size for each cell type is denoted in the label. Alveolar epithelial type 2, club, and differentiating basal cells are highlighted in blue. 10X: 10X Chromium.

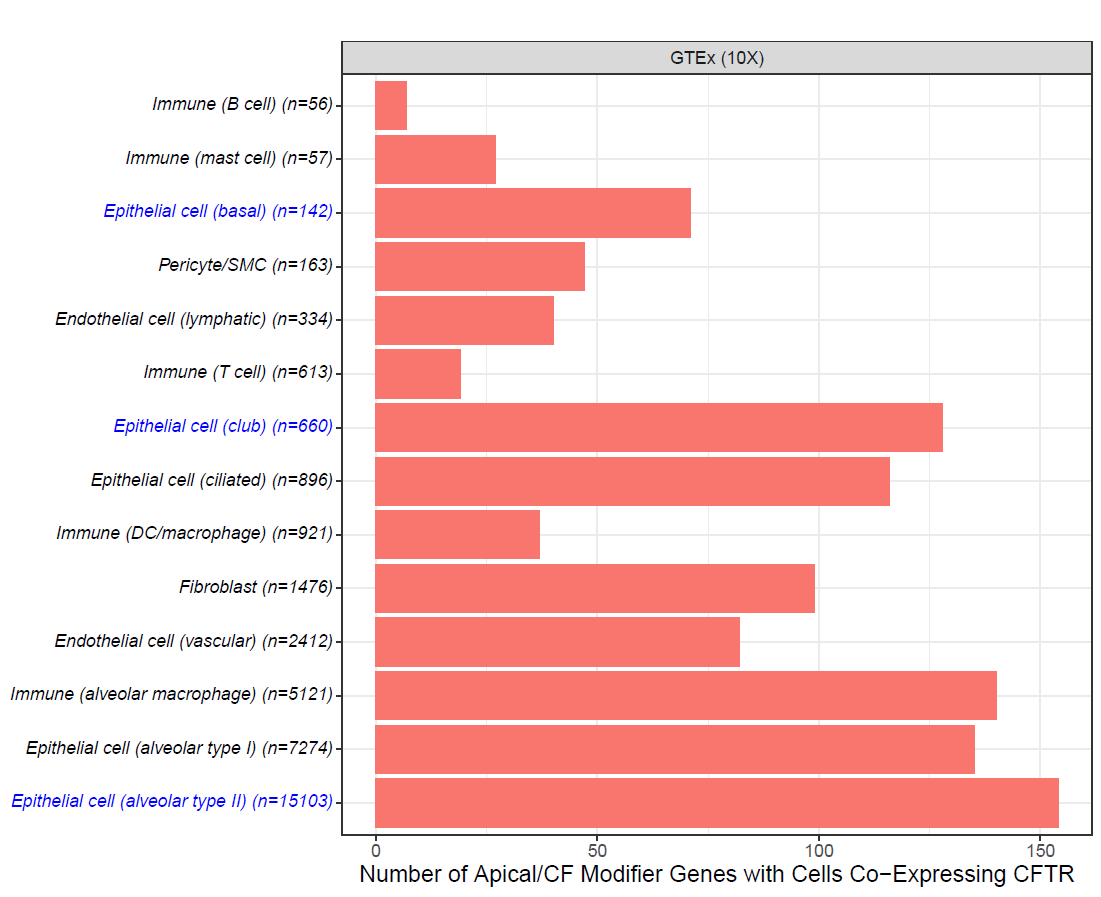

Figure S5. Number of cystic fibrosis modifier and apical constituent genes with at least one CFTR co-expressing cell across different lung cell type cells ordered by number of cells sequenced in the non-CF GTEx 10X Chromium data from GTEx ^23^.

A co-expressing cell was determined by observing detectable expression for CFTR and a given gene in the same cell. Sample size for each cell type is denoted in the label. Alveolar epithelial type 2, club, and basal cells are highlighted in blue. 10X: 10X Chromium.

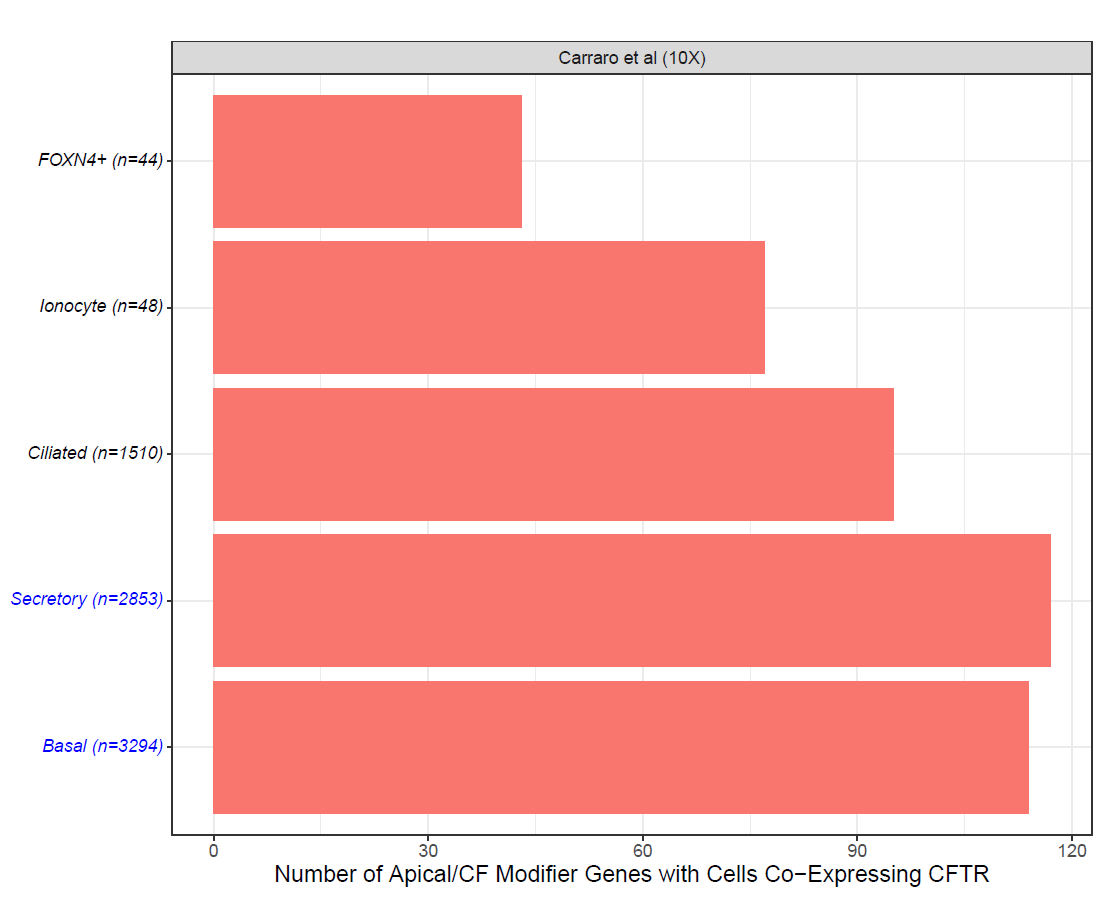

Figure S6. Number of cystic fibrosis modifier and apical constituent genes with at least one CFTR co-expressing cell across different lung cell type cells ordered by number of cells sequenced in the CF 10X Chromium data from Carraro et al. ^1^.

A co-expressing cell was determined by observing detectable expression for CFTR and a given gene in the same cell. Sample size for each cell type is denoted in the label. Secretory and basal cells are highlighted in blue. 10X: 10X Chromium.

#### 2.3 Lung cell types have different genome-wide co-expression profiles with CFTR

For non-CF samples in the SS2 data, AT2, club and differentiating basal cells have a high number of significant co-expressing relationships (Figure S7). Ionocytes showed no significant co-expressing genes due to its limited sample size of n=2.

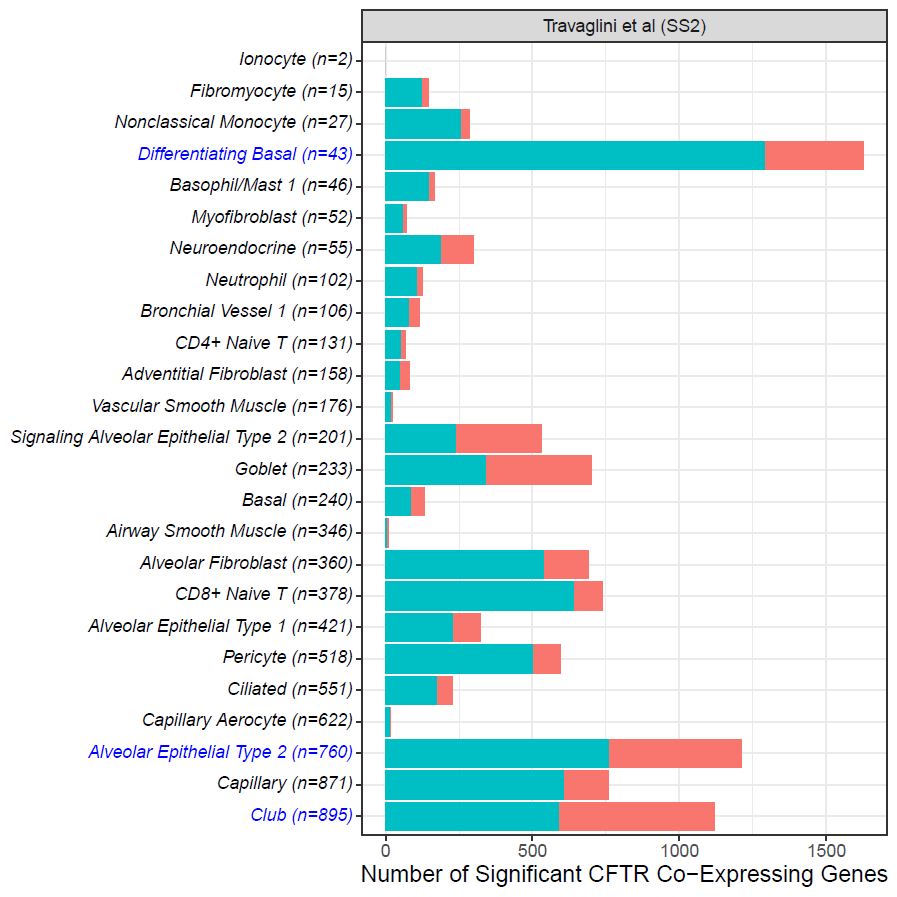

Figure S7. Number of significant co-expression relationships detected genome-wide (p < 0.05) across different lung cell types ordered by number of cells sequenced in the non-CF Smart-Seq2 dataset ^2^.

Contribution to the count of co-expression relationships for each cell type was based on the two models fitted for each gene, where each significant model result increased the count by 0.5. Number of cells sequenced for each cell type is denoted in the label. Counts of significant co-expressing genes are represented by the red bars, while counts of significant protein coding genes are shown by the teal bars. Alveolar epithelial type 2, club, and differentiating basal cells are highlighted in blue. SS2: Smart-Seq2.

In terms of functional themes among the co-expressing genes, Figure S8 shows cell type grouping when comparing associations with the top GO terms. Cell types with active CFTR co-expression networks including club, AT2 and differentiating basal cells appear to drive the GO term enrichment association results. These terms refer to important cellular functions including growth, development, biological regulation and response to external stimulus.

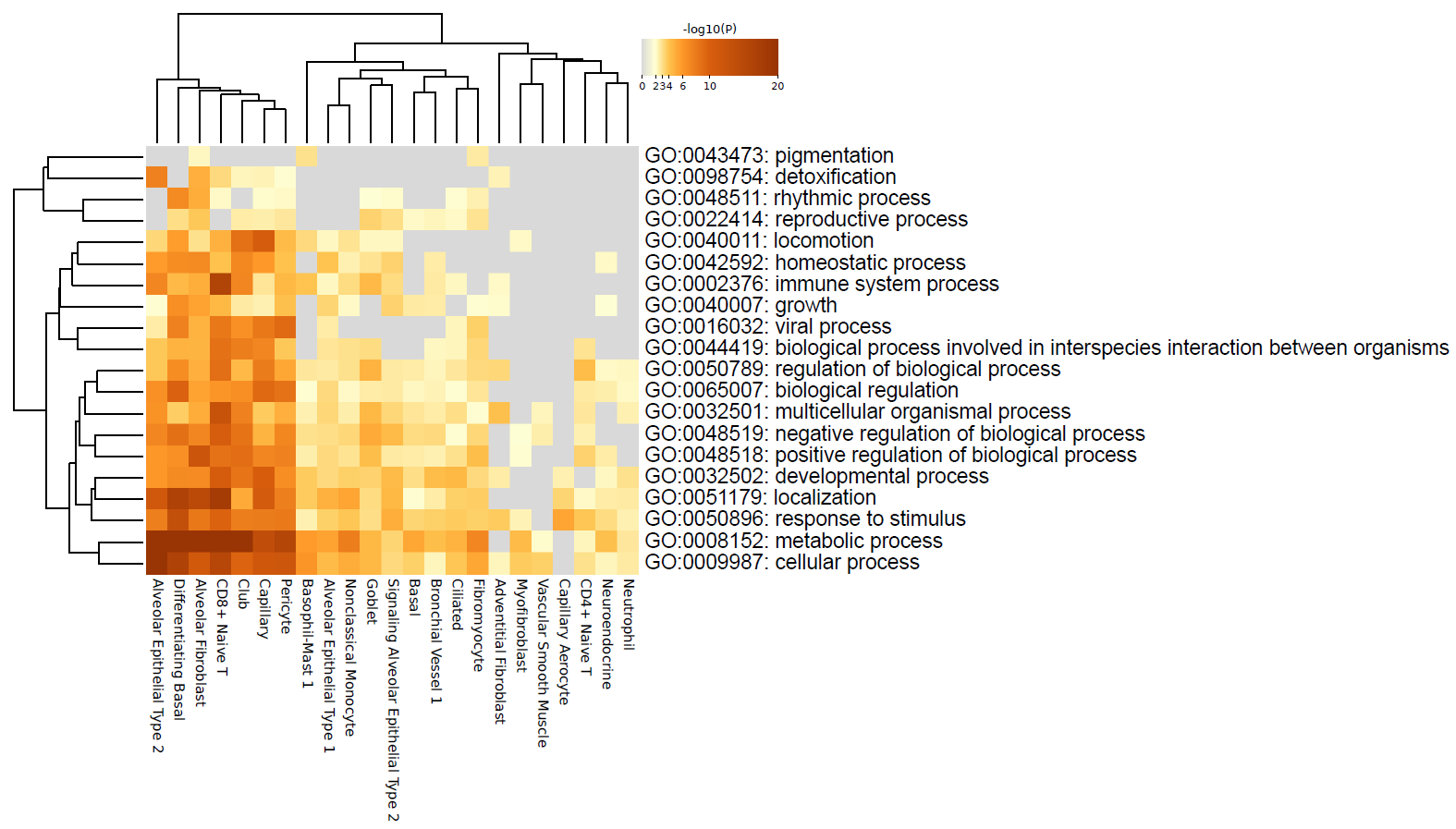

Figure S8. Functional GO term enrichment among the significant CFTR co-expressing genes in the non-CF Smart-Seq2 sample ^2^.

Heatmap displays the significance of association between a cell type’s co-expressing genes and a given GO term. Cell type (horizontal axis) and GO term (vertical axis) were clustered based on their enrichment results. Three groups of cell types in terms of differing GO term enrichment associations can be observed. The first group including club, AT2 and differentiating basal cells appear to drive the GO term enrichment association results. Cell types such as goblet cells formed the second group with less significant evidence of association with the same GO enrichment terms. The third group was composed of cell types with the fewest number of significant GO terms, like vascular smooth muscle cells.

As with the SS2 results, the 10X Chromium data also showed similar co-expression patterns among CF modifier and apical constituent genes (Figure S9, S10). In the non-CF 10X studies, both Travaglini et al. ^2^ and GTEx ^23^ 10X data showed high numbers of genes with at least one CFTR co-expressing differentiating basal, AT2 and club cell.

In the CF 10X sample, the broad basal cell group also had high numbers of co-expressing genes (Figure S11). Secretory cell types had the greatest number of co-expressions, though the difference with basal cells was small.

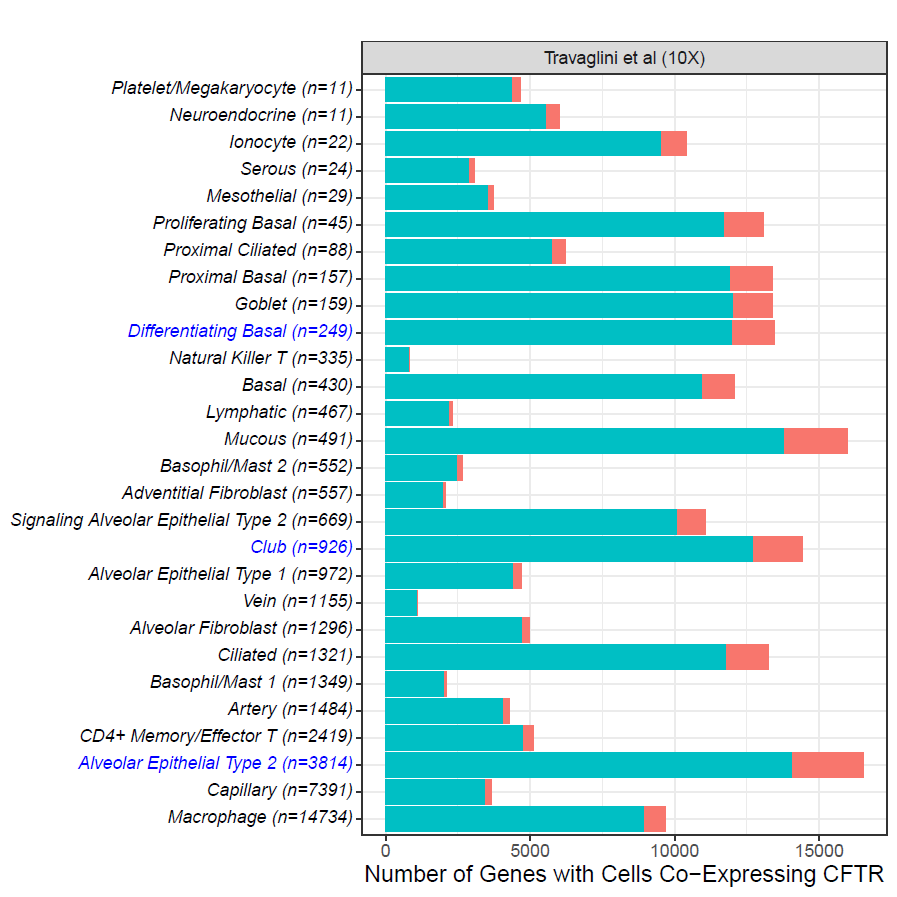

Figure S9. Number of genes with at least one CFTR co-expressing cell across different lung cell types ordered by number of cells sequenced in the non-CF 10X Chromium data from Travaglini et al. ^2^.

A co-expressing cell was determined by observing detectable expression for CFTR and a given gene in the same cell. Sample size for each cell type is denoted in the label. Counts of significant co-expressing genes are represented by the red bars, while counts of significant protein coding genes are shown by the teal bars. Alveolar epithelial type 2, club, and differentiating basal cells are highlighted in blue. 10X: 10X Chromium.

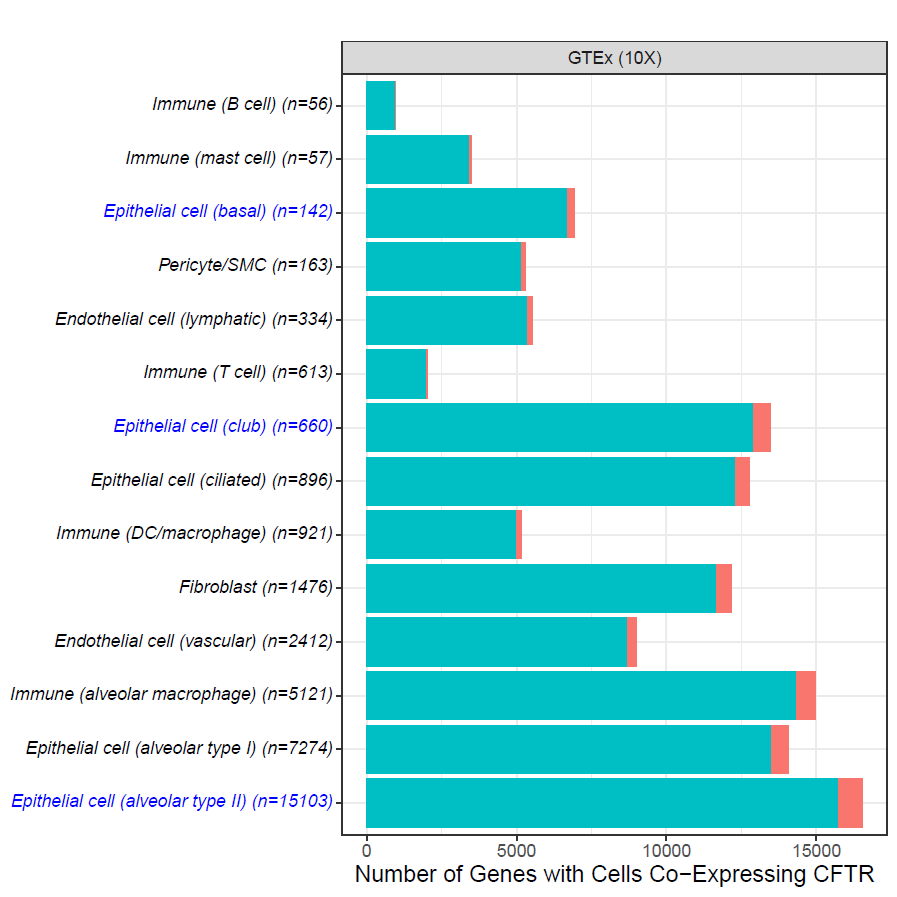

Figure S10. Number of genes with at least one CFTR co-expressing cell across different lung cell type cells ordered by number of cells sequenced in the non-CF GTEx 10X Chromium data from GTEx ^23^.

A co-expressing cell was determined by observing detectable expression for CFTR and a given gene in the same cell. Sample size for each cell type is denoted in the label. Counts of significant co-expressing genes are represented by the red bars, while counts of significant protein coding genes are shown by the teal bars. Alveolar epithelial type 2, club, and basal cells are highlighted in blue. 10X: 10X Chromium.

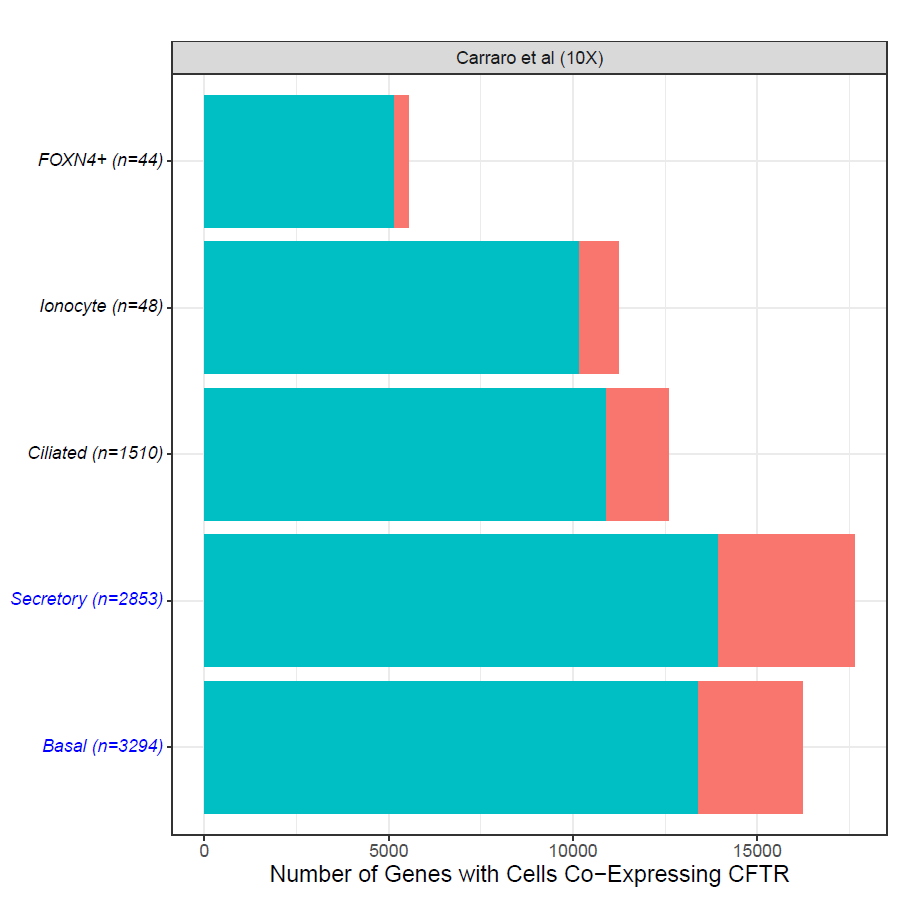

Figure S11. Number of genes with at least one CFTR co-expressing cell across different lung cell type cells ordered by number of cells sequenced in the CF 10X Chromium data from Carraro et al. ^1^.

A co-expressing cell was determined by observing detectable expression for CFTR and a given gene in the same cell. Sample size for each cell type is denoted in the label. Counts of significant co-expressing genes are represented by the red bars, while counts of significant protein coding genes are shown by the teal bars. Secretory and basal cells are highlighted in blue. 10X: 10X Chromium.

#### 2.4 Co-expression relationships among CFTR and CF lung modifier genes observed in alveolar epithelial type 2, differentiating basal and club cells

There is evidence of co-expression for CFTR-SLC9A3R2 in the differentiating basal cell as well as MUC4-SLC9A3R2 in the club cell (Figure S12). SLC9A3 encodes a Na+/H+ ion exchanger that has been associated with CF lung and intestine disease severity ^8,19^, and is down-regulated in CF primary human nasal epithelia cells upon infection with pseudomonas aeruginosa ^24^. The SLC9A3 protein complex which includes SLC9A3R2 is also shown to associate with CF lung disease ^21^. In addition to SLC9A3, the protein scaffold supported by SLC9A3R2 regulates CFTR and other ion transport pathways ^25^. The co-expression of CFTR and SLC9A3R2 in differentiating basal cells may represent the development of a coordinated ion transport scheme in the cell as it eventually differentiates into other cell types. On the other hand, MUC4 is another modifier for CF lung disease severity ^26^, and its gene product is a main component of airway mucus ^8^. Thus, co-expression of MUC4 and SLC9A3R2 in the club cells is consistent with its secretory functions in the lung.

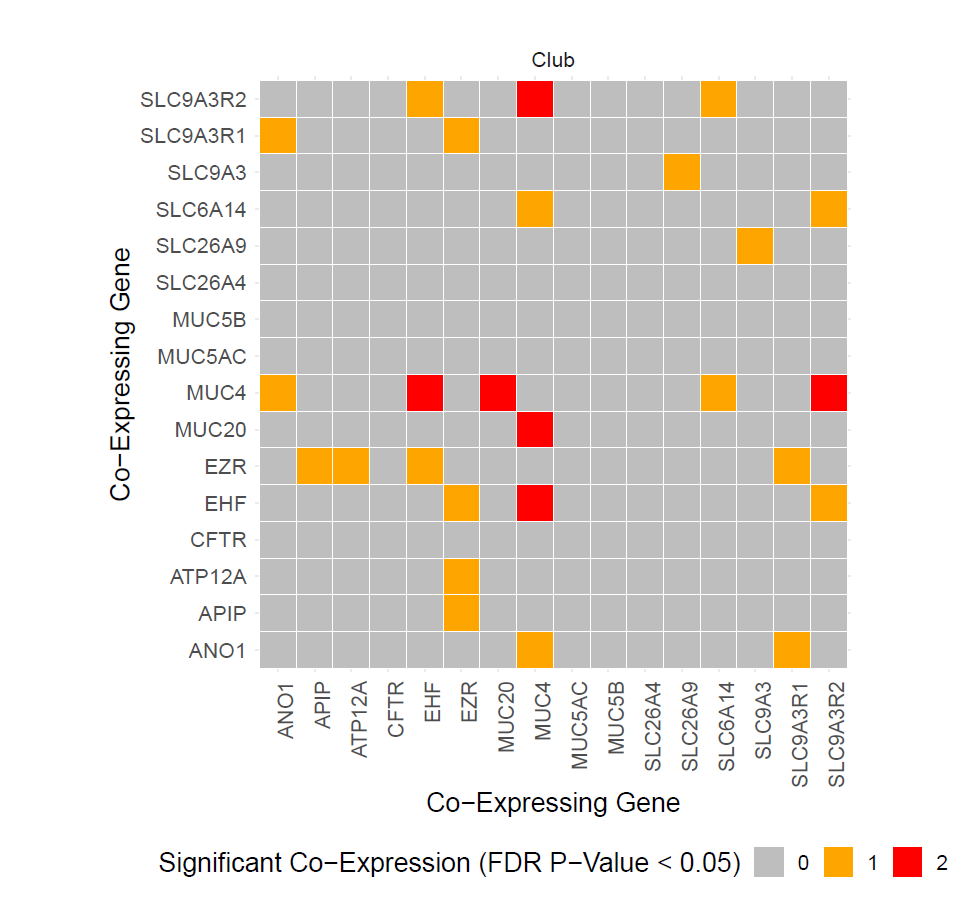

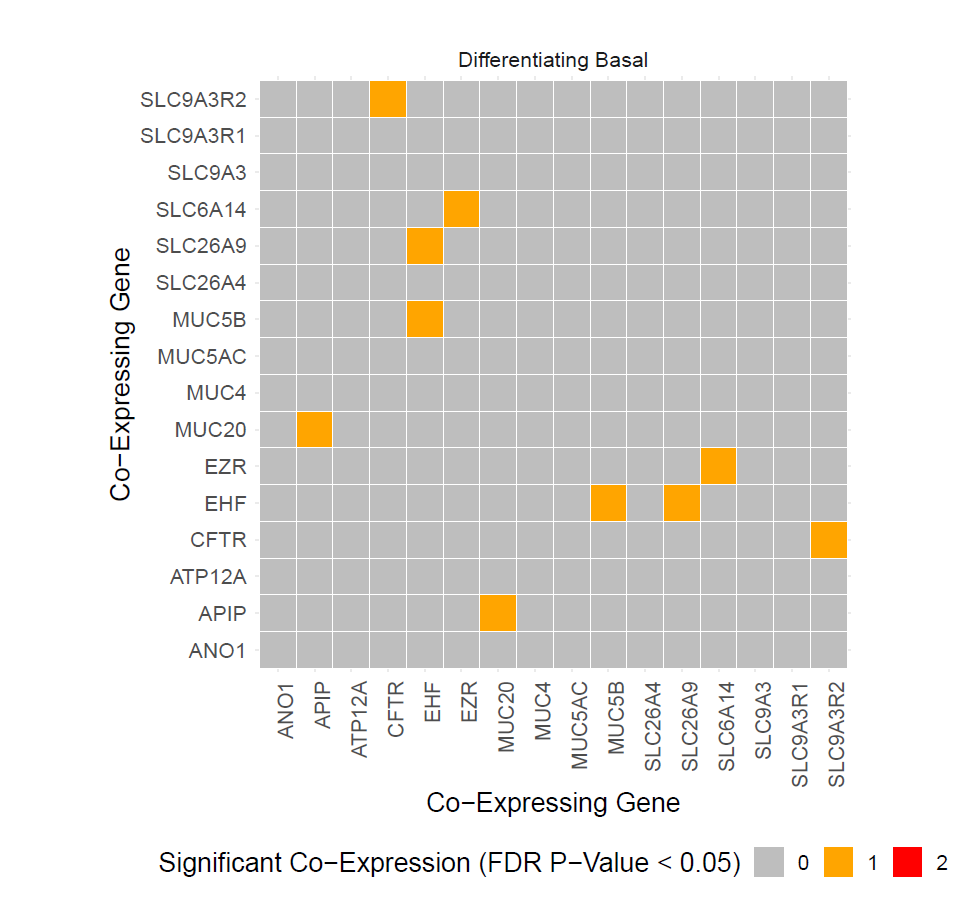

Figure S12. Co-expression relations among CFTR and cystic fibrosis modifier genes in club and differentiating basal cells from the non-CF Smart-Seq2 sample Travaglini et al. ^2^.

Significant co-expression relationships are shown in red (both co-expression model significant) or yellow (one significant co-expression model) for each pair of genes.

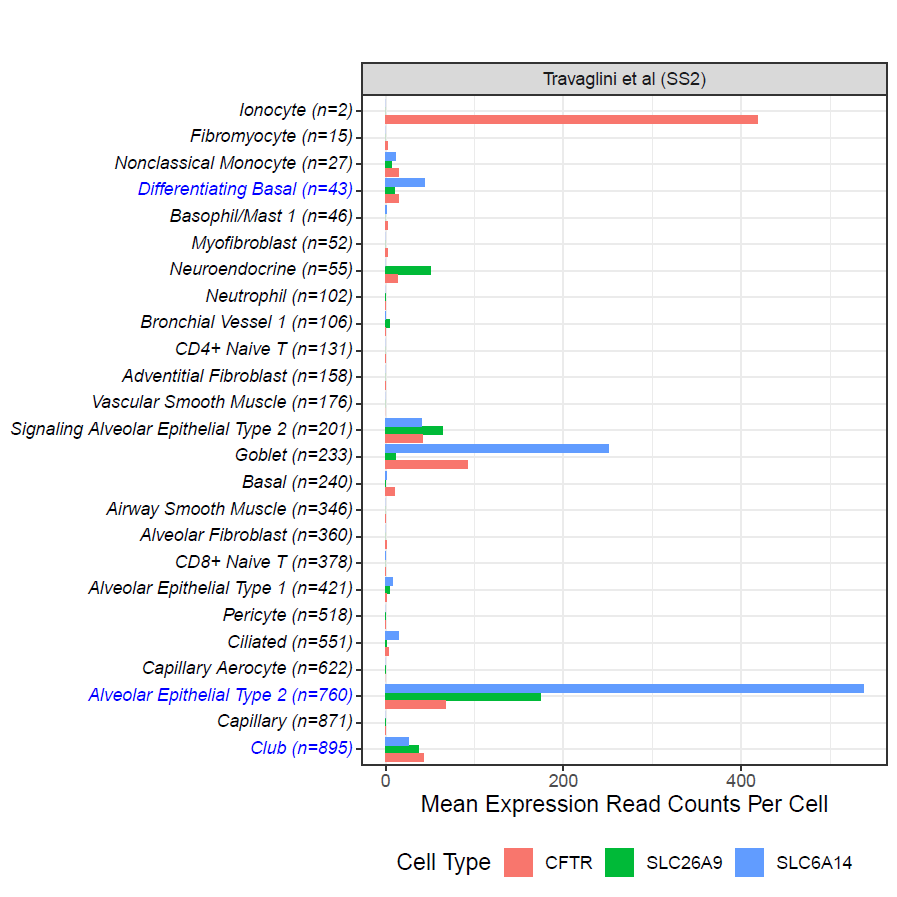

Figure S13. Expression levels of CFTR, SLC26A9 and SLC6A14 genes in different lung cell types from the non-CF Smart-Seq2 sample Travaglini et al. ^2^.

In Panel A, each bar for a given cell type represents the mean expression read counts per cell for either CFTR (red), SLC26A9 (green) and SLC6A14 (blue). Panel B shows the proportion of cells expressing CFTR, SLC26A9 and SLC6A14. Expressing cell was determined by observing detectable expression for CFTR (red), SLC26A9 (green) and SLC6A14 (blue) in the same cell. Cell types are arranged by their sample size in ascending order and denoted in the label. Alveolar epithelial type 2, club, and differentiating basal cells are highlighted in blue. SS2: Smart-Seq2.

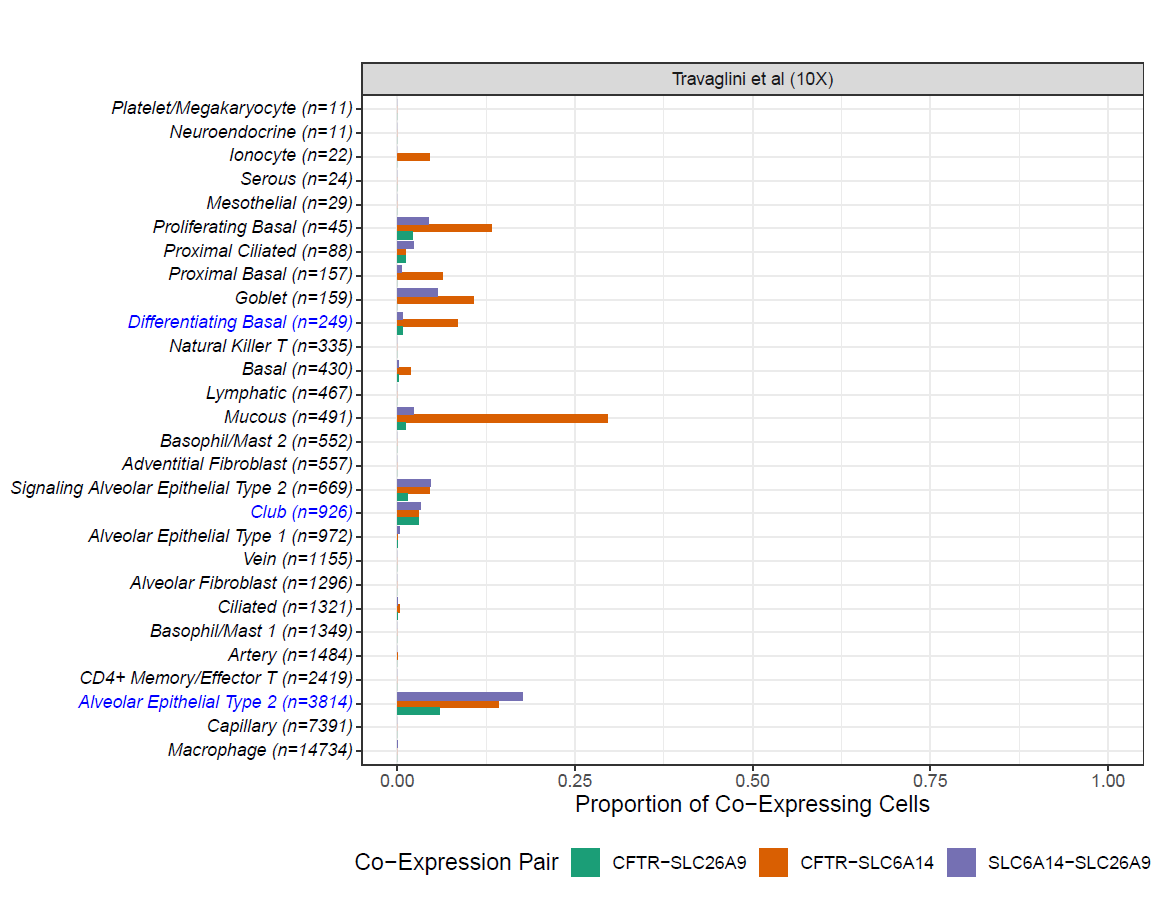

Figure S14. Proportion of cells co-expressing CFTR, SLC269 and SLC6A14 with each other in lung cell types of the non-CF 10X Chromium data from Travaglini et al. ^2^.

Co-expressing cell was determined by observing detectable expression for CFTR and SLC26A9 (green), CFTR and SLC6A14 (brown) and SLC6A14 and SLC26A9 (purple) in the same cell. Cell types are arranged by their sample size in ascending order and denoted in the label. Alveolar epithelial type 2, club, and differentiating basal cells are highlighted in blue. 10X: 10X Chromium.

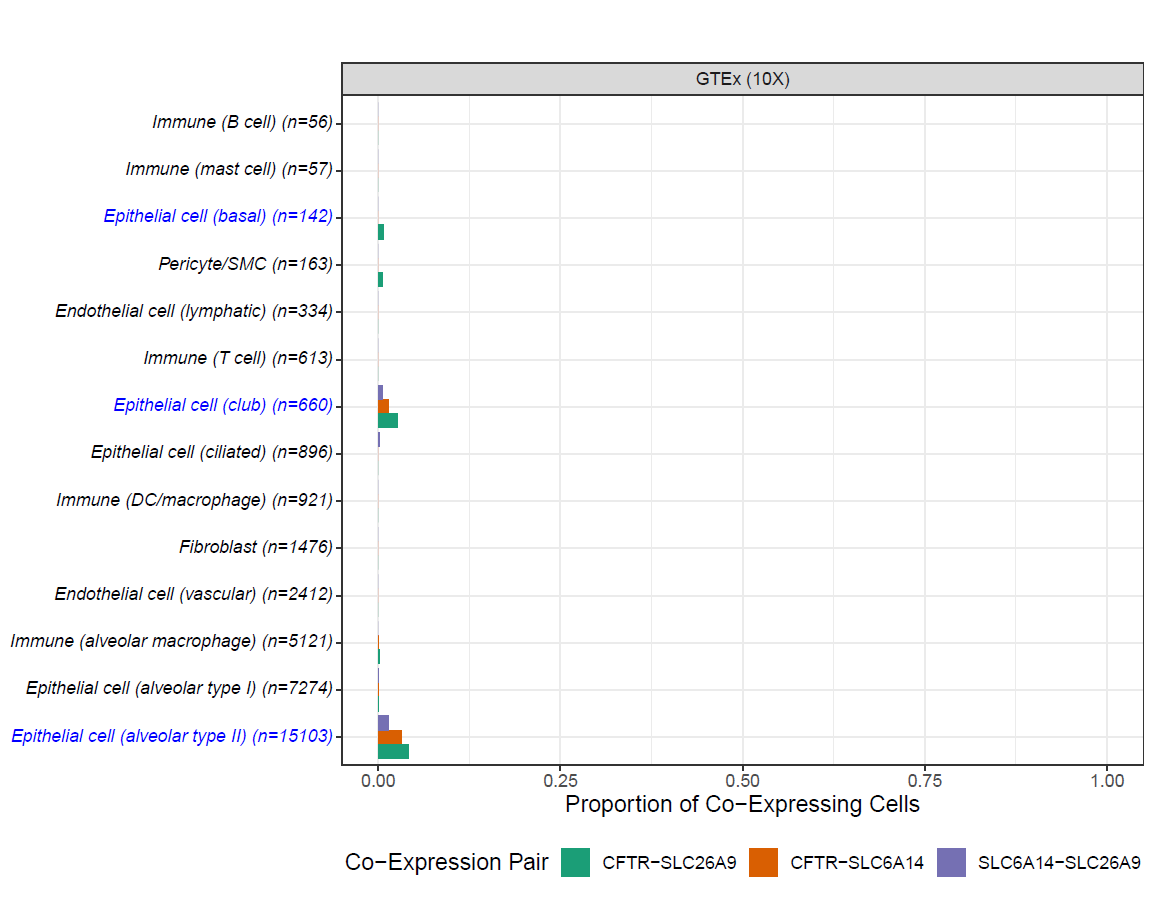

Figure S15. Proportion of cells co-expressing CFTR, SLC269 and SLC6A14 with each other in lung cell types of the non-CF GTEx 10X chromium data from GTEx ^23^.

Co-expressing cell was determined by observing detectable expression for CFTR and SLC26A9 (green), CFTR and SLC6A14 (brown) and SLC6A14 and SLC26A9 (purple) in the same cell. Cell types are arranged by their sample size in ascending order and denoted in the label. Alveolar epithelial type 2, club, and basal cells are highlighted in blue. 10X: 10X Chromium.

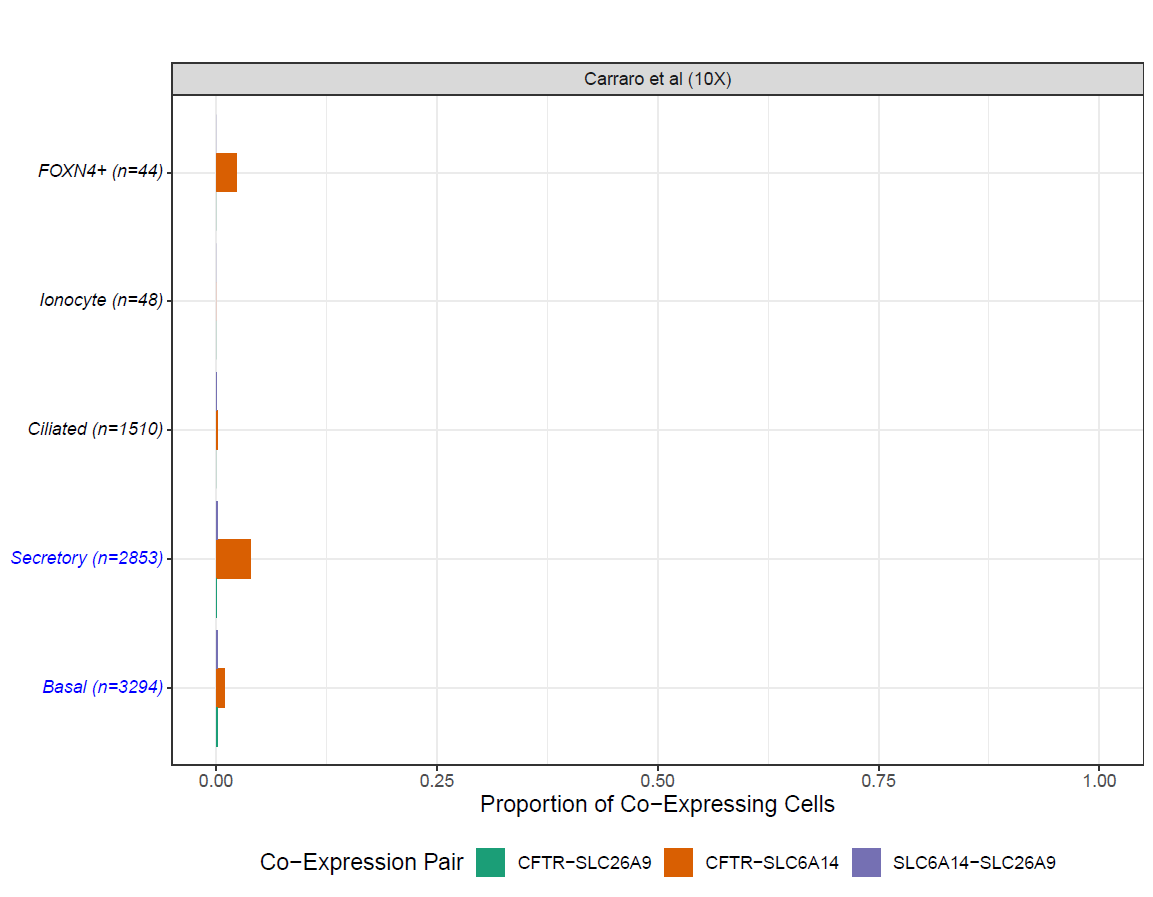

Figure S16. Proportion of cells co-expressing CFTR, SLC269 and SLC6A14 with each other in lung cell types of the CF 10X Chromium data from Carraro et al. ^1^.

Co-expressing cell was determined by observing detectable expression for CFTR and SLC26A9 (green), CFTR and SLC6A14 (brown) and SLC6A14 and SLC26A9 (purple) in the same cell. Cell types are arranged by their sample size in ascending order and denoted in the label. Secretory and basal cells are highlighted in blue. 10X: 10X Chromium.

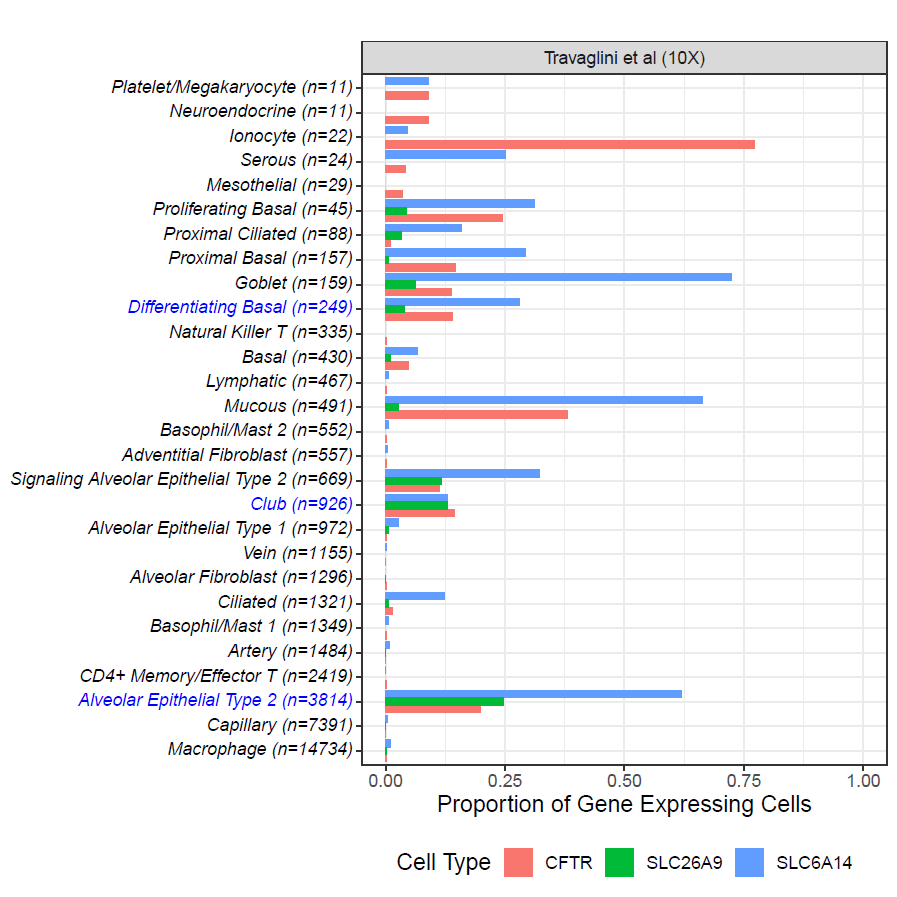

Figure S17. Proportion of cells expressing CFTR, SLC269 and SLC6A14 with each other in lung cell types of the non-CF 10X Chromium data from Travaglini et al. ^2^.

Expressing cell was determined by observing detectable expression for CFTR (red), SLC26A9 (green) and SLC6A14 (blue) in the same cell. Cell types are arranged by their sample size in ascending order and denoted in the label. Alveolar epithelial type 2, club, and differentiating basal cells are highlighted in blue. 10X: 10X Chromium.

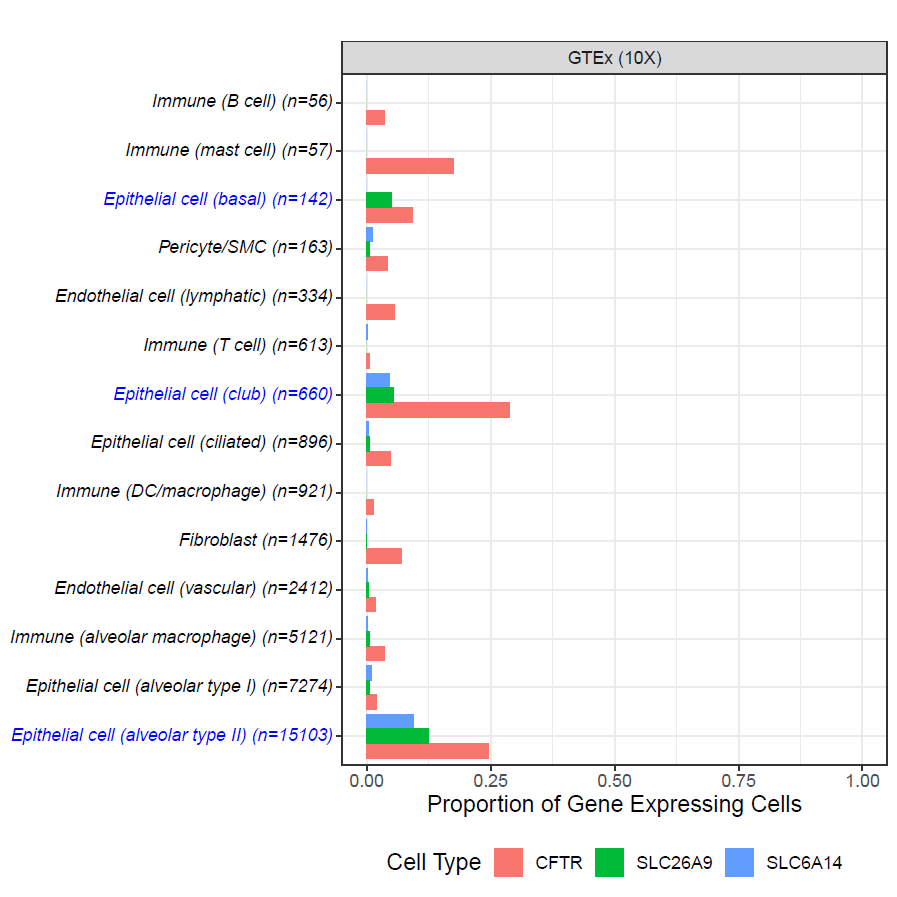

Figure S18. Proportion of cells expressing CFTR, SLC269 and SLC6A14 with each other in lung cell types of the non-CF GTEx 10X chromium data from GTEx ^23^.

Expressing cell was determined by observing detectable expression for CFTR (red), SLC26A9 (green) and SLC6A14 (blue) in the same cell. Cell types are arranged by their sample size in ascending order and denoted in the label. Alveolar epithelial type 2, club, and basal cells are highlighted in blue. 10X: 10X Chromium.

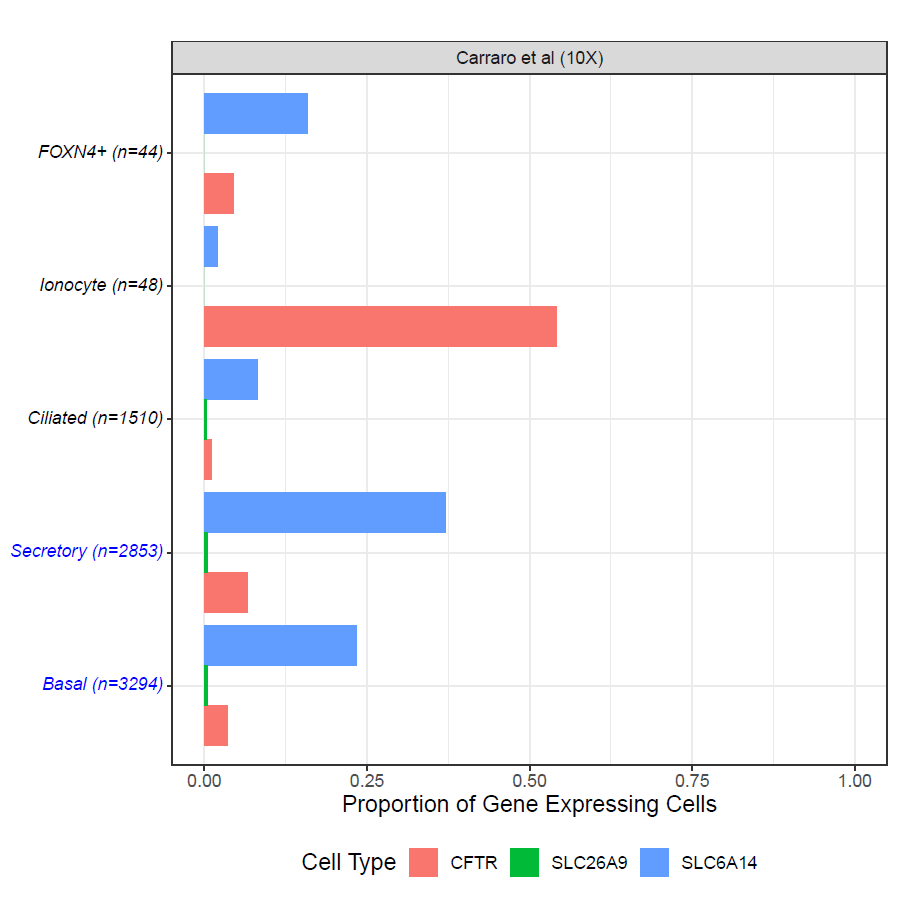

Figure S19. Proportion of cells expressing CFTR, SLC269 and SLC6A14 with each other in lung cell types of the CF 10X Chromium data from Carraro et al. ^1^.

Expressing cell was determined by observing detectable expression for CFTR (red), SLC26A9 (green) and SLC6A14 (blue) in the same cell. Cell types are arranged by their sample size in ascending order and denoted in the label. Secretory and basal cells are highlighted in blue. 10X: 10X Chromium.

Table S7. CFTR, SLC6A14 and SLC26A9 co-expression relations in other lung cell types from the non-CF Smart-Seq2 sample ^2^

| **Cell Type** | **Outcome Gene** | **Predictor Gene** | **Effect Size** | **P-Value** | **FDR P-Value** | **Cauchy Combination P-Value** |
| --- | --- | --- | --- | --- | --- | --- |
| Differentiating Basal | CFTR | SLC6A14 | -0.01968 | 0.28486 | 1 | 1 |
| Differentiating Basal | SLC6A14 | CFTR | -0.0014 | 0.59899 | 1 |  |
| Differentiating Basal | CFTR | SLC26A9 | -0.03189 | 0.18616 | 1 | 1 |
| Differentiating Basal | SLC26A9 | CFTR | -0.13948 | 0.19962 | 1 |  |
| Differentiating Basal | SLC6A14 | SLC26A9 | 0.00648 | 0.33729 | 1 | 1 |
| Differentiating Basal | SLC26A9 | SLC6A14 | -0.00306 | 0.09428 | 0.75424 |  |
| Club | CFTR | SLC6A14 | 0.00023 | 0.34401 | 1 | 1 |
| Club | SLC6A14 | CFTR | 0.00106 | 0.40467 | 1 |  |
| Club | CFTR | SLC26A9 | 3.36x10^-05^ | 0.9798 | 1 | 1 |
| Club | SLC26A9 | CFTR | -0.00039 | 0.58713 | 1 |  |
| Club | SLC6A14 | SLC26A9 | 0.00088 | 0.47992 | 1 | 1 |
| Club | SLC26A9 | SLC6A14 | 0.00115 | 0.23309 | 1 |  |
| Goblet | CFTR | SLC6A14 | 0.00012 | 0.03193 | 0.15422 | 0.3299 |
| Goblet | SLC6A14 | CFTR | 0.0007 | 0.26442 | 0.69785 |  |
| Goblet | CFTR | SLC26A9 | -0.00334 | 0.21371 | 0.61472 | 0.61472 |
| Goblet | SLC26A9 | CFTR | 2.0057320NA | NA | NA |  |
| Goblet | SLC6A14 | SLC26A9 | 0.00871 | 0.00903 | 0.06081 | 0.06081 |
| Goblet | SLC26A9 | SLC6A14 | NA | NA | NA |  |
| Signaling Alveolar Epithelial Type 2 | CFTR | SLC6A14 | 0.00099 | 0.68686 | 1 | 1 |
| Signaling Alveolar Epithelial Type 2 | SLC6A14 | CFTR | -0.00012 | 0.93178 | 1 |  |
| Signaling Alveolar Epithelial Type 2 | CFTR | SLC26A9 | 0.00221 | 0.07401 | 0.73431 | 0.22152 |
| Signaling Alveolar Epithelial Type 2 | SLC26A9 | CFTR | 0.00284 | 0.00329 | 0.09365 |  |
| Signaling Alveolar Epithelial Type 2 | SLC6A14 | SLC26A9 | 0.00268 | 0.22358 | 1 | 1 |
| Signaling Alveolar Epithelial Type 2 | SLC26A9 | SLC6A14 | 0.00036 | 0.44986 | 1 |  |
